# A PATIENT-DERIVED EX VIVO TISSUE MODEL OF CHOLANGIOCARCINOMA USING PRECISION-CUT TISSUE SLICES

**DOI:** 10.64898/2026.09.17.745977

**Authors:** Timothy M Gilbert, Owen McGreevy, Maria-Danae Jessel, Roz Jenkins, Mohamed Bosakhar, Marc Quinn, Lawrence O’Leary, Anthony Evans, Timothy Andrews, Rafael Diaz-Nieto, Robert P Jones, Stephen Fenwick, William Greenhalf, Daniel Palmer, Hassan Z Malik, Christopher Goldring, Laura Randle

## Abstract

**Background & Aims:** Cholangiocarcinoma (CCA) is an aggressive malignancy with poor five-year survival. Improving outcomes requires preclinical models that faithfully recapitulate the native tumour microenvironment. Precision-Cut Tissue Slices (PCTS) retain in-vivo architecture and represent a promising candidate. We aimed to establish a patient-derived PCTS model of CCA, characterise its response to ex-vivo culture, and assess its use as a platform for therapeutic testing.

**Methods:** PCTS were generated from 25 patients undergoing curative-intent resection for presumed CCA (2022–2024) and cultured for up to 15 days. Viability and histological architecture were assessed serially. Quantitative proteomics (SWATH-DIA) was performed at Days 0, 3, 7 and 15. A subset were treated with staurosporine or clinically relevant chemotherapy (5′-deoxy-5-fluorouridine, gemcitabine ± cisplatin).

**Results:** PCTS viability was maintained to Day 15, with no significant reduction from baseline. PCTS retain tumour architecture, cytokeratin-19-positivity and resident CD3+/CD68+ immune cells during culture. Proteomic profiling quantified 4,578 proteins and identified a staged response, with differentially abundant proteins increasing from 8 (Day 3) to 127 (Day 7) and 180 (Day 15), stabilising thereafter. This comprised of an early loss of inflammatory and stromal proteins, sustained cellular stress responses and metabolic reprogramming. CCA subtype-specific proteomic signatures were preserved throughout culture. Staurosporine produced dose-dependent cytotoxicity whilst chemotherapy responses were variable.

**Conclusions:** Patient-derived CCA PCTS maintain viability, tumour architecture and subtype identity for 15 days while undergoing a defined proteomic response to culture and retain pharmacological responsiveness.

**GRAPHICAL ABSTRACT:** 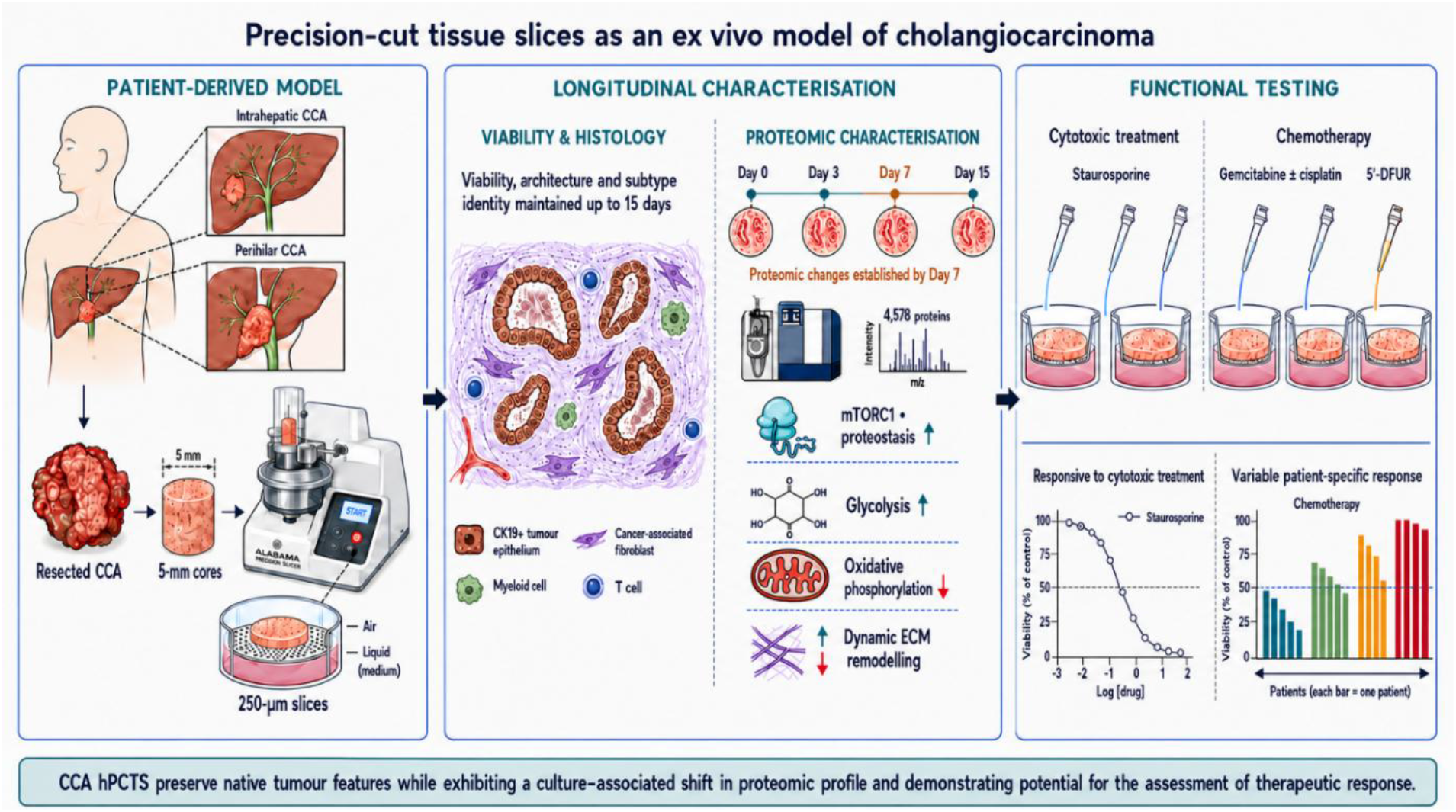

**IMPACT AND IMPLICATIONS:** PCTS are a potentially valuable preclinical cancer model, retaining the native TME, while offering a 3Rs approach to translational research, yet their use in CCA remains limited and the model has not been comprehensively characterised. This study establishes patient-derived CCA PCTS as a viable ex-vivo platform and provides the first detailed proteomic characterisation of how this model behaves during prolonged culture. Our proteomic findings are informative more broadly for researchers developing PCTS as a platform in other cancer types, underscoring the impact of ex-vivo culture and the need to consider whether the proteomic changes described here may influence model behaviour and potential drug sensitivity.

## 1.0 INTRODUCTION

Cholangiocarcinoma (CCA) is an aggressive gastrointestinal malignancy with rising global incidence ^1^. Surgical resection remains the only curative option but is available to a minority of patients as most present with advanced disease ^2,3^. Outcomes are further limited by a propensity for chemoresistance and high rates of relapse after resection and adjuvant treatment ^4–7^. Consequently, overall five-year survival rates remain low, between 5-10%^8,9^. There is an urgent need to improve understanding of this difficult-to-treat disease and identify novel therapeutic strategies.

One of the major hurdles in furthering our understanding of tumour biology and the subsequent development of new treatments is the ability to produce pre-clinical models that faithfully recapitulate the *in vivo* cancer^10^. This is highlighted by the high rates of attrition seen during the transition of novel therapeutics into phase I oncology trials. An analysis of clinical trial data from 2010 to 2017 shows the primary reason for failure of anti-cancer drugs in early phase trials were a lack of clinical efficacy (40%– 50%), unmanageable toxicity (30%) and poor drug-like properties (10%–15%)^11^. Since a significant proportion of drug development fails due to a lack of clinical efficacy, an increasing emphasis has been placed on the development of preclinical models to help evaluate and rationalise potential new treatments^11^. However, developing a pre-clinical model that encapsulates all the complexities and nuances of an *in vivo* tumour is exceptionally challenging.

Cellular cultures and murine models remain the most used techniques to explore CCA biology, each with their own advantages and disadvantages^12^. Human precision-cut tissue slice (PCTS) models have re-emerged as a potentially valuable tool for studying human cancer biology. Unlike 2D and 3D *in-vitro* models that can oversimplify intricate cancer-stromal interactions, PCTS retain the complex architecture and native tumour microenvironment of the original tissue they derive from ^13^. This theoretically provides researchers with a more representative model, as well as a more clinically relevant platform for therapeutic studies^13^.

A significant advantage of PCTS is that it is a relatively low cost and easily adaptable technique that can be utilised on diseased and healthy tissue from multiple different sites (lung, liver, prostate, pancreas, brain etc.)^13–17^. PCTS also align with the 3Rs principles of replacement, reduction and refinement, offering a route to reduce reliance on animal models in cancer research. In 2010, de Graaf *et al.* published a protocol which became the reference method for researchers working with tissue slice cultures and helped to standardize techniques leading to a renewed interest in the method^18^. More recently this technique has been successfully adapted to generate PCTS from primary and metastatic liver tumours with variable survivorship^19^. PCTS have been successfully used to model drug response and resistance across a range of different cancer types^20,21^. Despite this, it remains an underutilised technique in the study of CCA^22^.

Here we report our experience in the development of PCTS from surgically resected specimens of CCA and offer an in-depth proteomic characterisation of this model and assess its use as a platform for therapeutic intervention.

## 2.0 MATERIAL AND METHODS

### 2.1 TISSUE COLLECTION

All patients undergoing surgical resection of a presumed CCA were eligible for tissue donation. Written informed consent was obtained from each patient in accordance with Human Tissue Authority (HTA) guidelines, and the study was conducted under the PINCER platform study (REC ref: 15/NW/0477) in accordance with the Declaration of Helsinki ^23^. Following surgical excision, a section of tumour tissue was removed under histopathology supervision to preserve resection-margin assessment, collected immediately into ice-cold Belzer University of Wisconsin (UW)® Cold Storage Transplant Solution (Bridge to Life, Columbia, SC, USA), and transferred to the Human Liver Research Facility (University of Liverpool) for processing (Figure 1).

**Figure 1.**
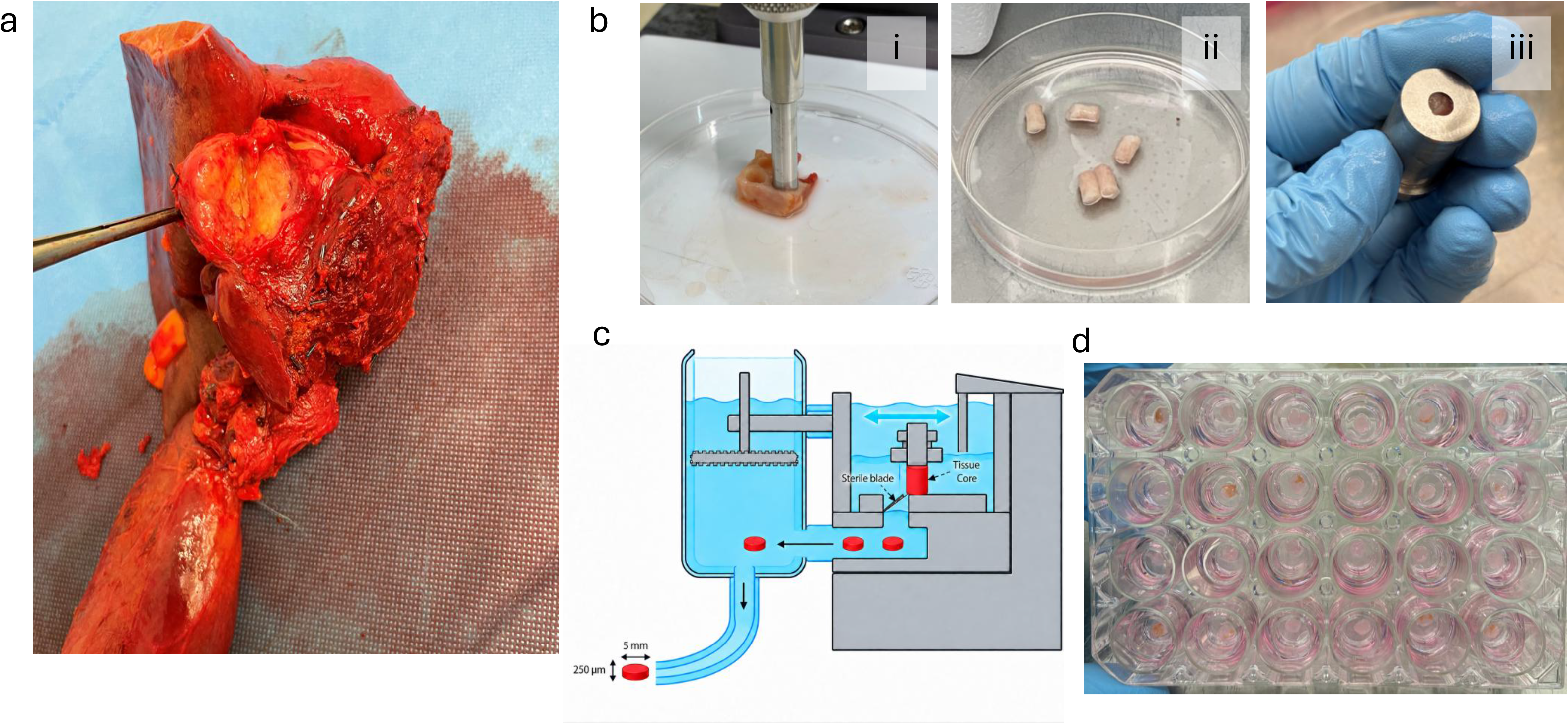
Resected cholangiocarcinoma can be reproducibly processed into precision-cut tissue slices for culture. (A) Representative resected cholangiocarcinoma specimen opened for tumour sampling. (B) Generation of a tumour core and loading into the tissue holder of a Krumdieck live tissue microtome. (C) Schematic of precision-cut tissue slicing to generate slices 250 µm thick and 5 mm in diameter under chilled conditions. (D) Representative 24-well culture plate containing cholangiocarcinoma tissue slices on Millicell® organotypic inserts. CCA, cholangiocarcinoma; hPCTS, human precision-cut tissue slices.

### 2.2 GENERATION OF CCA PCTS

CCA PCTS were generated based on previously published protocols ^18,24^. Tumour specimens were transferred to the Human Liver Research Facility and processed using a 5 mm coring drill (Alabama Research and Development, Munford, AL, USA) to produce multiple cylindrical tissue cores. Cores were sliced on a Krumdieck Tissue Slicer (Alabama Research and Development) in 1× Krebs-Henseleit buffer (Supplementary Table 1) at 4 °C to generate 250 µm slices (arm speed 4, blade speed 4; Figure 1). Slices were collected from the glass trap of the slicer and recovered in supplemented Williams’ E medium (WEM; Supplementary Table 2) for 1 h at 37 °C before culture.

### 2.3 EX-VIVO CULTURE OF CCA PCTS

Following recovery, PCTS were transferred to fresh 24-well plates on 12 mm, 0.4 µm Millicell® organotypic inserts (Merck) and cultured on an orbital shaker at 95 rpm under normoxic conditions (37 °C, 5% CO_2_). Each well contained 450 µL supplemented WEM with 200 ng/mL human epidermal growth factor (hEGF; Corning). Media were changed on Day 1 and every 48 h thereafter, for as long as required.

### 2.4 MTS VIABILITY ASSAY

Tissue slice viability was assessed using the CellTiter 96® AQueous One Solution Cell Proliferation Assay (MTS; Promega) and absorbance read at 490 nm on a Varioskan Flash microplate reader (Thermo Fisher Scientific).

### 2.5 PCTS THERAPEUTIC DOSING

CCA PCTS were maintained overnight for recovery and treated from Day 1 for 48 h with staurosporine (STS), gemcitabine with or without cisplatin (GemCis), or 5′-deoxy-5-fluorouridine (DFUR), with a complete medium and drug exchange at 24 h. Each agent was tested across a half-log concentration series with matched vehicle controls (0.5% dimethyl sulfoxide [DMSO] and 0.9% saline) and untreated controls; STS served as a positive control in the chemotherapy experiments. GemCis was co-administered at a fixed gemcitabine:cisplatin ratio of 40:1. Three technical replicates were performed per condition, with viability subsequently analysed at patient level. Following treatment, PCTS were collected for viability and downstream analyses.

### 2.6 FFPE BLOCK PREPARATION, MICROTOMY & SECTIONING

Tissue slices fixed in 10% neutral-buffered formalin were processed to formalin-fixed paraffin-embedded (FFPE) blocks and cut into 4 µm sections. Sections were stained with haematoxylin and eosin (H&E) or used for immunohistochemistry (IHC). IHC was performed for cleaved caspase-3 (CC3), Ki67, cytokeratin-19 (CK19), CD3 and CD68, with heat-induced antigen retrieval on a Dako PT-Link and detection by horseradish peroxidase (HRP)-conjugated secondary antibody and diaminobenzidine (DAB). All slides for a given marker were stained within a single batch and processed together.

Stained slides were scanned on an Aperio CS2 slide scanner at 20× magnification and analysed in QuPath v0.6.0. A single set of colour deconvolution stain vectors was estimated from representative tissue and applied across all slides. Positive cells were detected using the Positive Cell Detection function, with single-intensity thresholds set with reference to matched positive and negative control tissue and applied consistently across the slide set. Total and positive cell counts were recorded to calculate percentage positivity.

### 2.7 PROTEOMIC SAMPLE PREPARATION AND SWATH-MS ACQUISITION

#### 2.7.1 Spectral library preparation

A pooled spectral library was generated from bulk resection tissue from nine CCA patients (5 iCCA and 4 pCCA), five of whom also contributed PCTS to the SWATH-DIA sample set. Total protein (1.5 mg) was reduced, alkylated and digested in solution using a Trypsin/Lys-C Mix, prefractionated by strong cation exchange (SCX) chromatography, desalted, and analysed by data-dependent acquisition (DDA).

#### 2.7.2 CCA tumour PCTS sample preparation (SWATH-DIA)

Protein was extracted from individual tumour PCTS by homogenisation in urea lysis buffer, and 100 µg per sample was reduced, alkylated and digested with trypsin using single-pot solid-phase-enhanced sample preparation (SP3).

#### 2.7.3 LC–MS and data acquisition (library fractions and SWATH samples)

Peptides were separated on a bioZen XB-C18 column (75 µm × 250 mm) and analysed on a TripleTOF 6600 mass spectrometer (SCIEX). Spectral library fractions were acquired by DDA and individual PCTS samples by SWATH-DIA using 100 variable windows across 400–1500 m/z. The library was searched in ProteinPilot 5.1 (Paragon algorithm) against the UniProt Swiss-Prot human database at 1% false discovery rate (FDR) and exported using PeakView. The PeakView-exported spectral library and matched UniProt Swiss-Prot FASTA were subsequently used for DIA-NN processing of the SWATH-DIA data.

#### 2.7.4 DIA-NN processing and quantification

The individual PCTS SWATH-DIA samples were processed using DIA-NN (data-independent acquisition neural networks; v1.8)^25^, with the project-specific spectral library and the matched UniProt Swiss-Prot FASTA. Processing used double-pass library generation with match-between-runs enabled, Trypsin/P specificity with one missed cleavage, and fixed carbamidomethylation. Peptide and protein identifications were controlled at 1% FDR and quantification performed at protein-group level.

## 3.0 BIOINFORMATIC & STATISTICAL ANALYSIS

### 3.1 CLINICAL AND ROUTINE LABORATORY STATISTICAL ANALYSIS

Analyses were performed in SPSS Statistics v22 (IBM), GraphPad Prism v10.4 (GraphPad Software) and R v4.4.2. Normality was assessed with the Shapiro-Wilk test. Normally distributed variables are reported as mean ± SD and compared with two-sided Student’s t-tests; non-normal data are reported as median (interquartile range) and compared with non-parametric tests. Metabolic activity and immunohistochemical marker positivity were assessed across culture timepoints as repeated measures within patient, using mixed-effects models or the Friedman test as appropriate, with comparisons against Day 0. Baseline (Day 0) viability was compared between iCCA and pCCA by Welch’s two-sided t-test. Significance was set at p ≤ 0.05 unless otherwise stated.

### 3.2 PROTEOMIC ANALYSIS

SWATH-DIA proteomic data were analysed in R v4.4.2. Protein-group intensities from DIA-NN were log2-transformed and missing values imputed by random forest (imp4p) with CCA subtype as the conditioning variable. Differential abundance across culture timepoints (Day 0, 3, 7 and 15) was modelled in limma using an additive design (Batch + Subtype + Day), with within-donor correlation accounted for by duplicateCorrelation and empirical Bayes moderation; all pairwise contrasts among Days 0, 3, 7 and 15 were tested, with proteins considered differentially abundant at Benjamini–Hochberg FDR ≤ 0.05 and |log2 fold-change| ≥ 0.5. Differences between anatomical subtypes were assessed using an interaction design (Batch + Day x Subtype) with the same donor blocking and moderation, taking the subtype coefficient at the reference day as the Day 0 contrast, and whether the temporal response differed by subtype was tested by a joint moderated F test across the three Day-by-subtype interaction coefficients. Because imputation was conditioned on subtype, the Day 0 subtype contrast was restricted before model fitting to protein groups with sufficient measured values in each subtype, leaving 3,750 of 4,578 (Supplementary Methods). Functional pathway enrichment was assessed by gene set enrichment analysis against the MSigDB Hallmark collection (fgsea). Temporal protein trajectories were clustered by fuzzy c-means (Mfuzz) on the overall cohort, with subtype-specific mean trajectories overlaid on the shared clusters and cluster-level enrichment assessed against Reactome (ReactomePA).

## 4.0 RESULTS

### 4.1 PATIENT DEMOGRAPHICS

Twenty-five patients who underwent curative-intent resection for presumed CCA between 2022 and 2024 (median age 71 [IQR 62-75]; 52% male) were included in this study (Supplementary Table 3). Most patients (n=17) underwent an anatomical right or left hepatectomy with or without radical bile duct resection. All bulk tumour specimens from which our PCTS were derived underwent a full histopathological assessment at the Royal Liverpool University Hospital. Twenty-three patients were confirmed to have biliary tract cancer (12 iCCA, 11 pCCA), two presumed iCCA cases were subsequently reclassified as hepatocellular carcinoma and squamous cell carcinoma (Supplementary Table 3).

### 4.2 EX-VIVO VIABILITY

CCA PCTS maintained viability over a 15-day culture period. Mean MTS absorbance increased from 1.56 (95% CI 1.32–2.16) at Day 0 to 1.78 (95% CI 1.37–2.61) at Day 7. Thereafter, viability reduced by an average of 40% relative to day 7. Mean MTS absorbance did not differ significantly from Day 0 at Day 11 (p = 0.14) or Day 15 (p = 0.21; Figure 2). The proportion of CC3+ and Ki67+ cells remained stable throughout culture, with no significant differences from Day 0 (Figure 2). At Day 0, MTS absorbance was higher in iCCA than pCCA tissue slices (p = 0.048; Supplementary Table 4). Tumour cellularity was also higher in iCCA, with a median of 4,932 CK19-positive cells per 4 µm section compared with 701 in pCCA, which may contribute to this difference.

**Figure 2.**
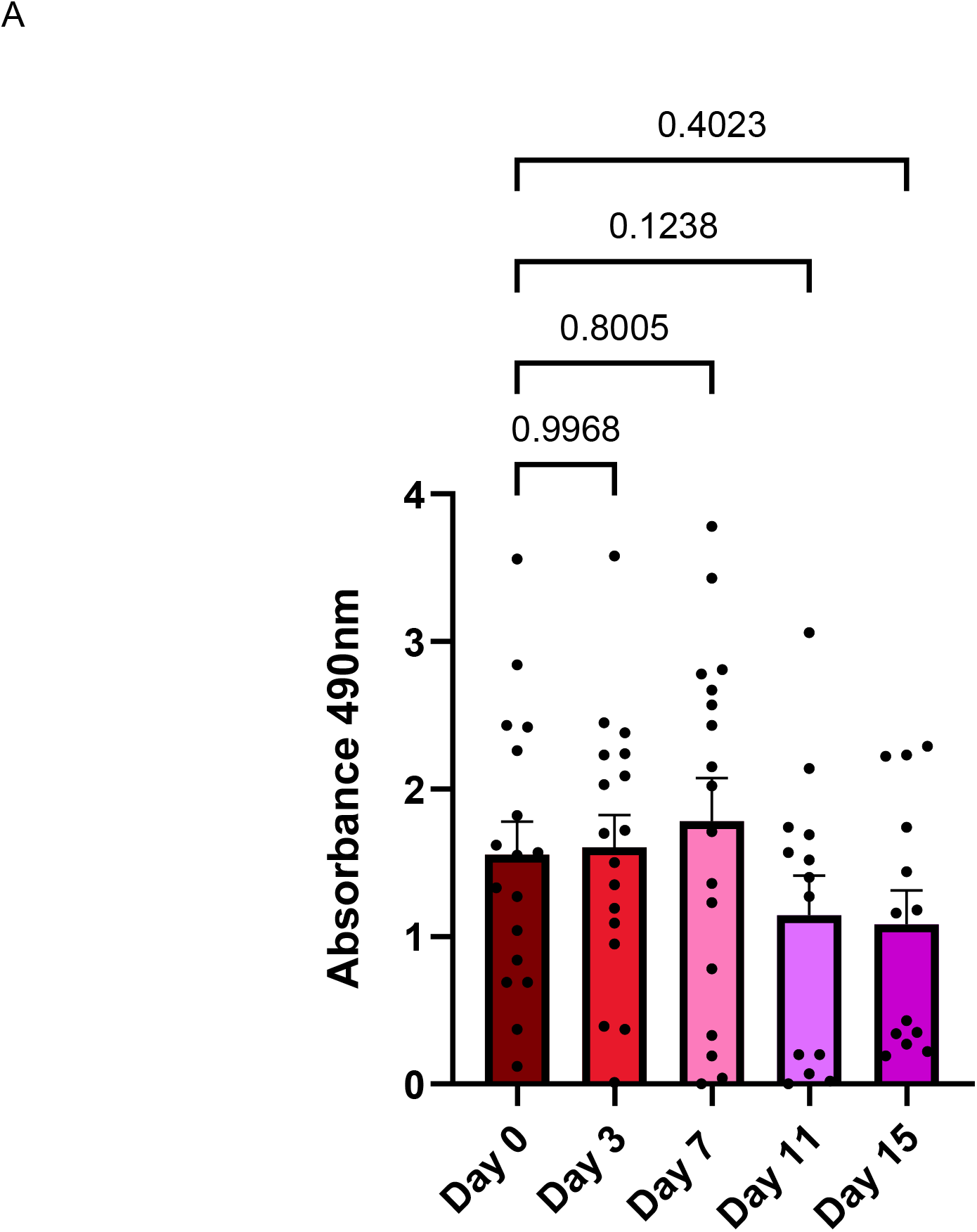
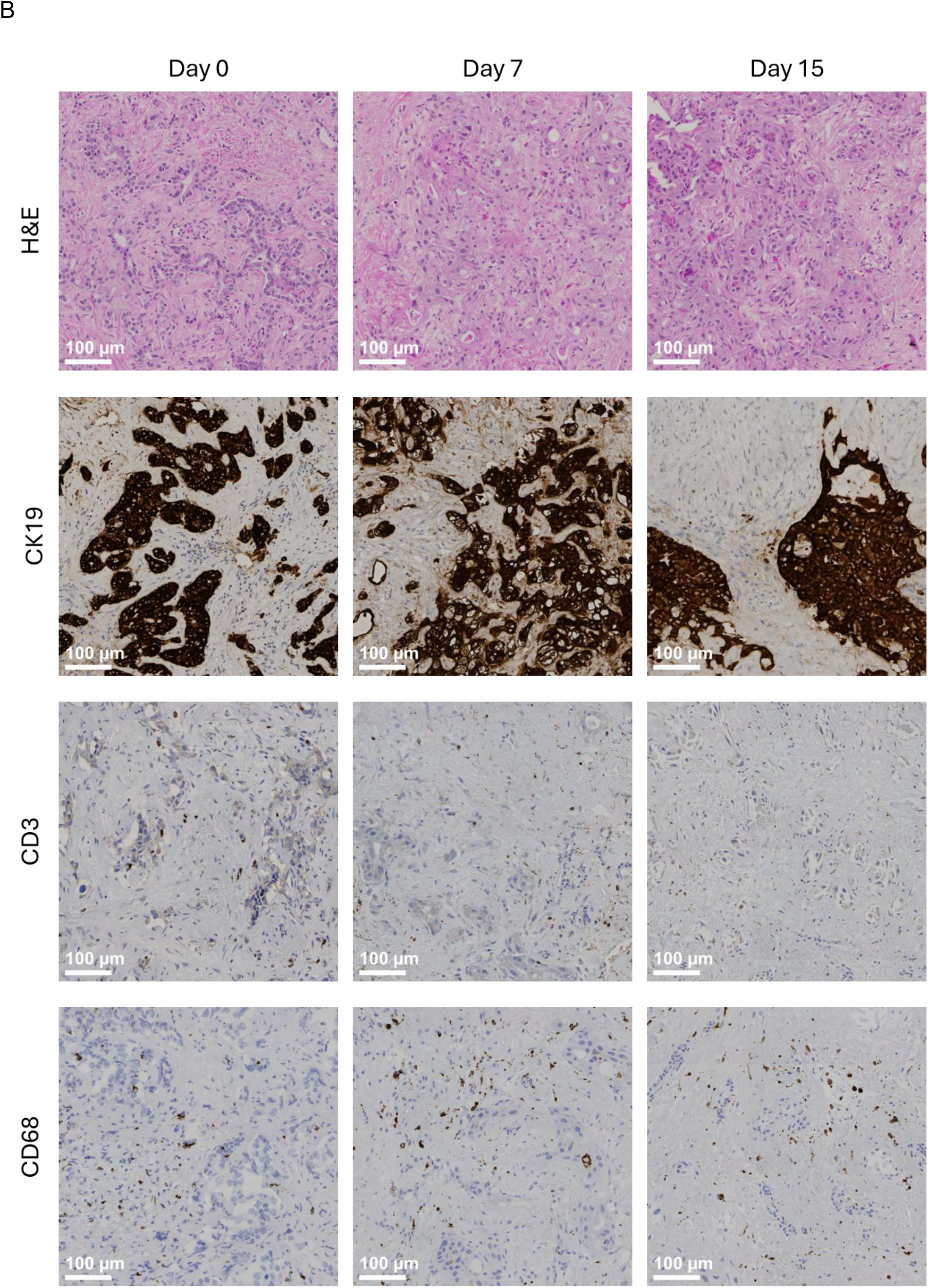
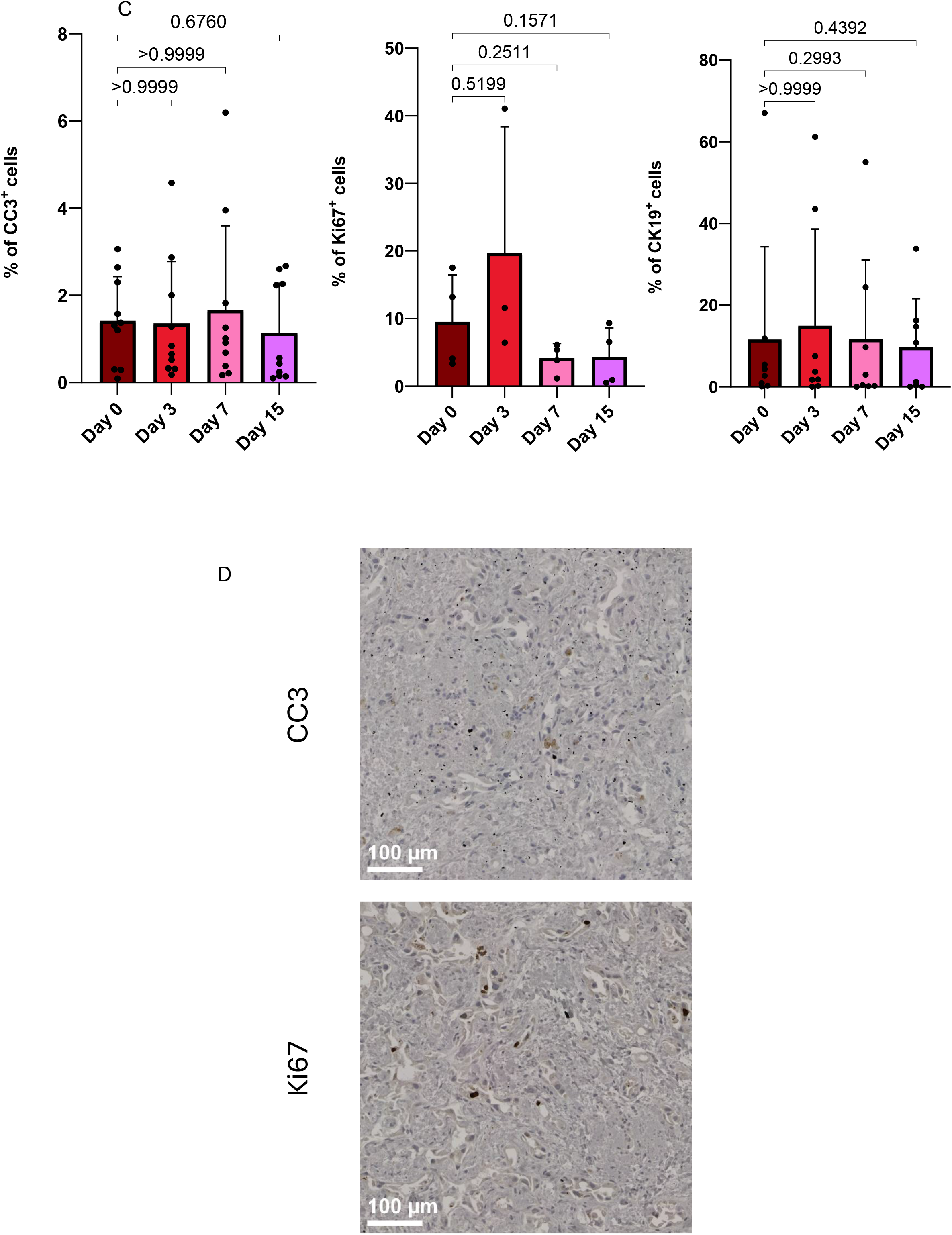
Cholangiocarcinoma tissue slices remain viable and retain tumour architecture across 15 days of culture. (A) Tissue slice viability across Days 0, 3, 7, 11 and 15 assessed by MTS assay at 490 nm. Each point represents one patient, calculated from the mean of technical replicates; n = 17 at Days 0, 3 and 7 and n = 13 at Days 11 and 15. Bars show mean ± SD. Comparisons with Day 0 were performed using a mixed-effects model with Dunnett’s multiple-comparisons test. (B) Representative H&E, CK19, CD3 and CD68 staining at Days 0, 7 and 15. (C) CC3 (n = 13), Ki67 (n = 4) and CK19 (n = 13) positivity across culture, expressed as the percentage of positive cells per slice. Bars show mean ± SD. CK19 and CC3 were analysed using Friedman tests with Dunn’s multiple comparisons against Day 0; Ki67 was analysed using a mixed-effects model with Dunnett’s multiple comparisons against Day 0. (D) Representative CC3-positive apoptotic cells and Ki67-positive proliferating cells at Day 15. Scale bars, 100 µm. CC3, cleaved caspase-3; CK19, cytokeratin 19; H&E, haematoxylin and eosin; hPCTS, human precision-cut tissue slices; MTS, 3-(4,5-dimethylthiazol-2-yl)-5-(3-carboxymethoxyphenyl)-2-(4-sulfophenyl)-2H-tetrazolium.

### 4.3 HISTOLOGICAL PHENOTYPE

Macroscopically all specimens required a tumour mass that could provide a sample large enough to produce the cores required for slicing. As a result, all the pCCA tumours included in this study demonstrated a macroscopic mixed growth pattern (Mass Forming + Periductal infiltrating), with a central mass that could be sampled at the liver hilum. iCCA samples were also of a mass forming phenotype but peripherally located.

Microscopically PCTS retain the key morphological features of the source tumour throughout culture, including widely spaced CK19+ tubulo-glandular epithelial structures surrounded by dense desmoplastic stroma. CD3+ and CD68+ cells were identified within cultured slices, indicating retention of resident immune populations (Figure 2).

### 4.4 PROTEOMIC CHARACTERISATION OF CCA PCTS

To characterise the response of CCA-PCTS to ex-vivo culture, longitudinal proteomic profiling was performed at Day 0, 3, 7 and 15. In total, 4,578 proteins were quantified across all samples (1% FDR). Differential abundance analysis identified 8, 127 and 180 proteins that differed from Day 0 at Days 3, 7 and 15, respectively (Figure 3), indicating that the principal proteomic transition occurred during the first week of culture. Integration of differential abundance analysis, GSEA and Mfuzz temporal clustering identified several key biological responses (Figures 3 and 4).

**Figure 3.**
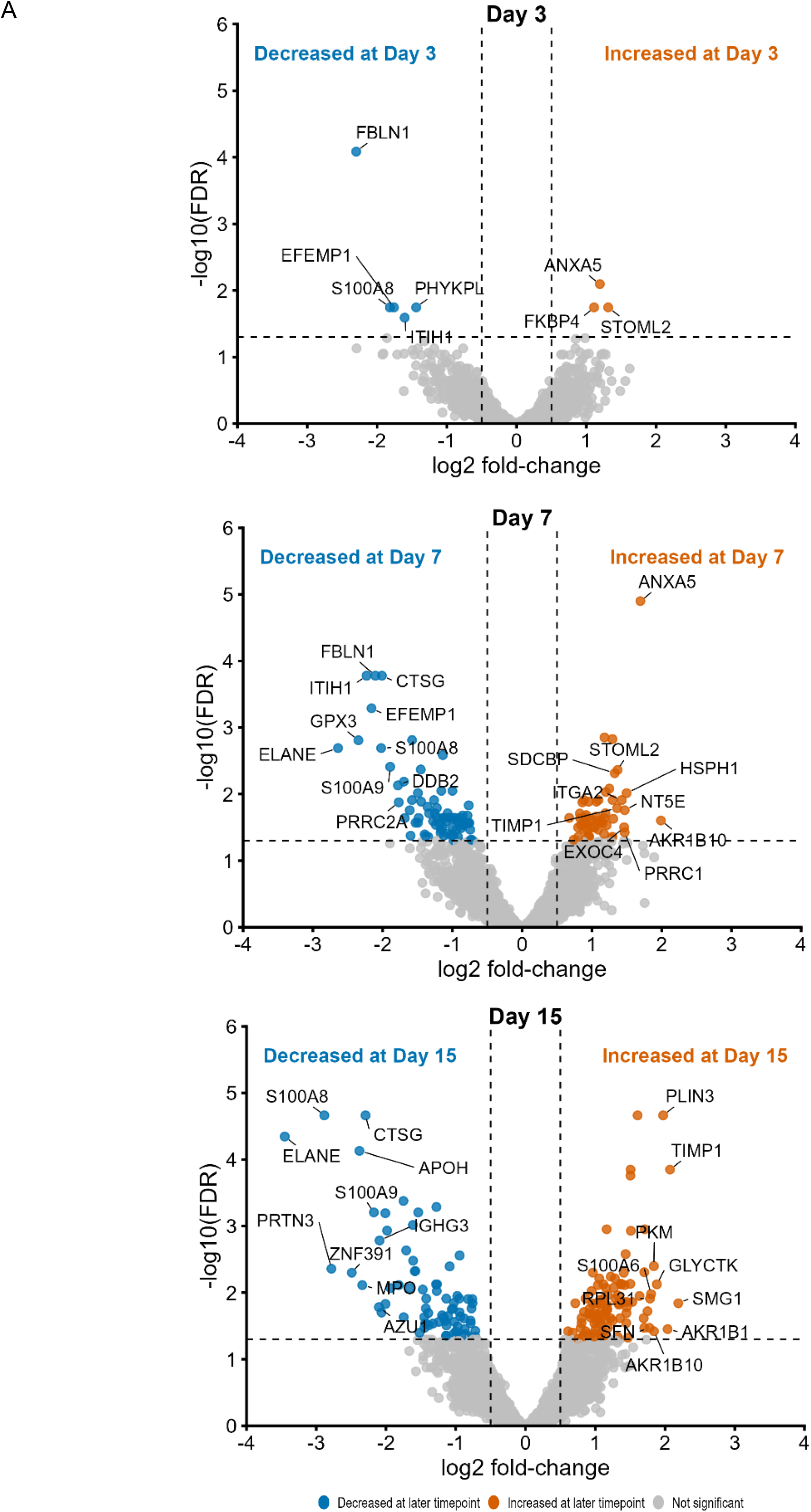
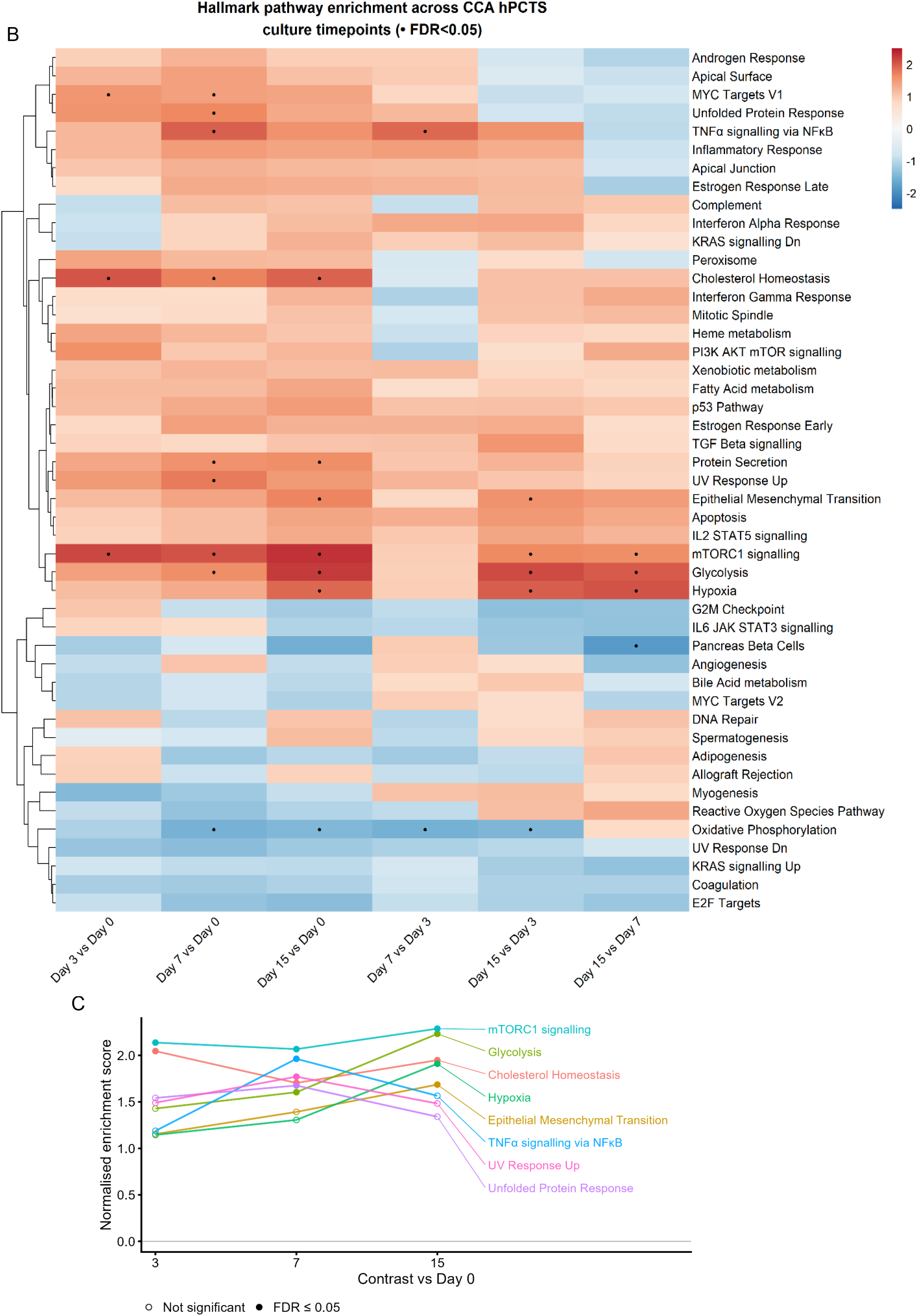
Cholangiocarcinoma tissue slices show selective proteomic and pathway remodelling during culture. **(A)** Volcano plots showing differential protein abundance at Days 3, 7 and 15 relative to Day 0. Orange indicates increased abundance, blue decreased abundance and grey non-significant proteins; differential abundance was defined as FDR ≤ 0.05 and |log2 fold-change| ≥ 0.5. (B) Hallmark pathway enrichment across temporal contrasts. Colour represents normalised enrichment score (NES), with red indicating positive and blue negative enrichment; points indicate FDR ≤ 0.05. (C) NES trajectories for selected Hallmark pathways at Days 3, 7 and 15 relative to Day 0. Filled points indicate FDR ≤ 0.05 and open points indicate non-significant enrichment. FDR, false discovery rate; NES, normalised enrichment score.

**Figure 4.**
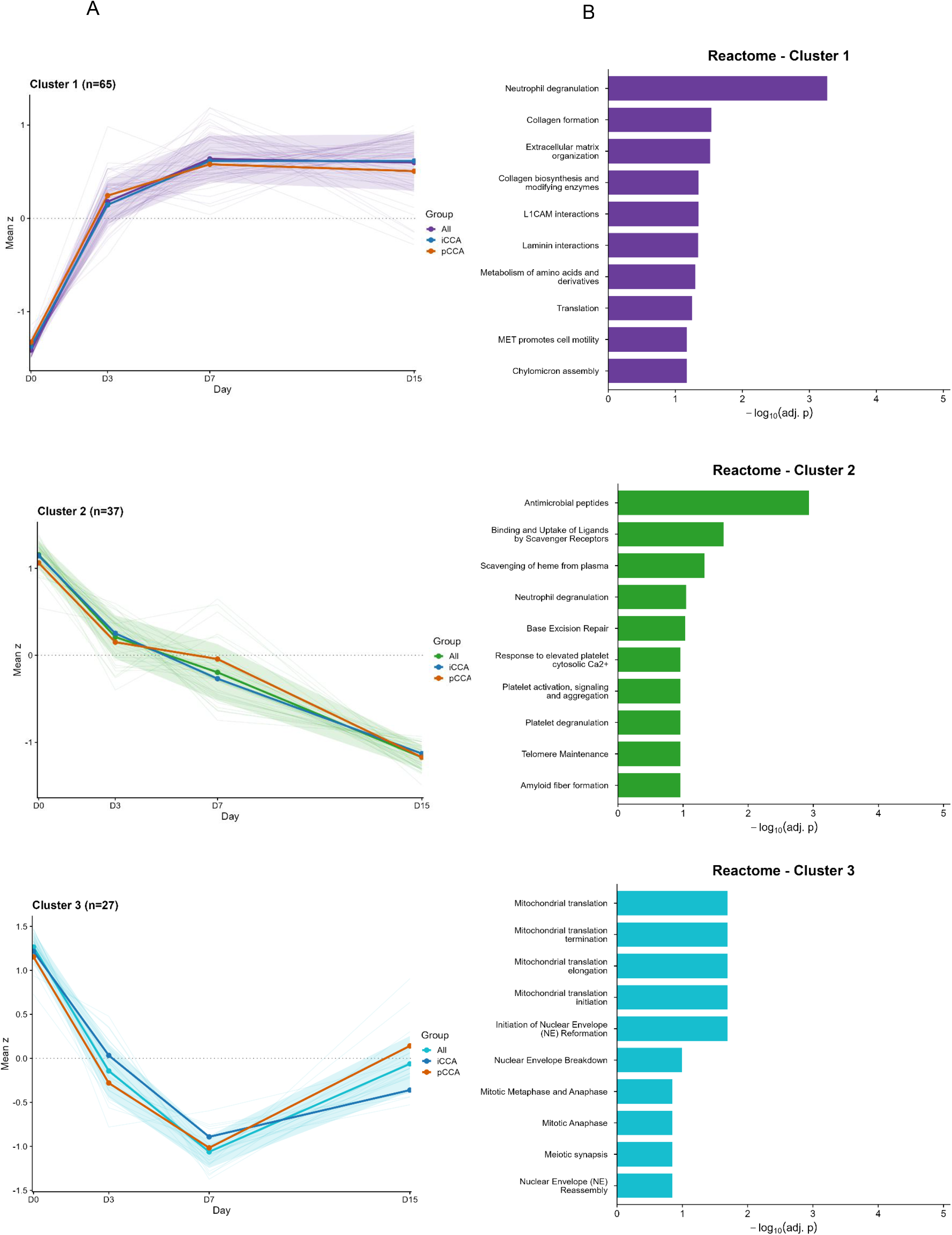
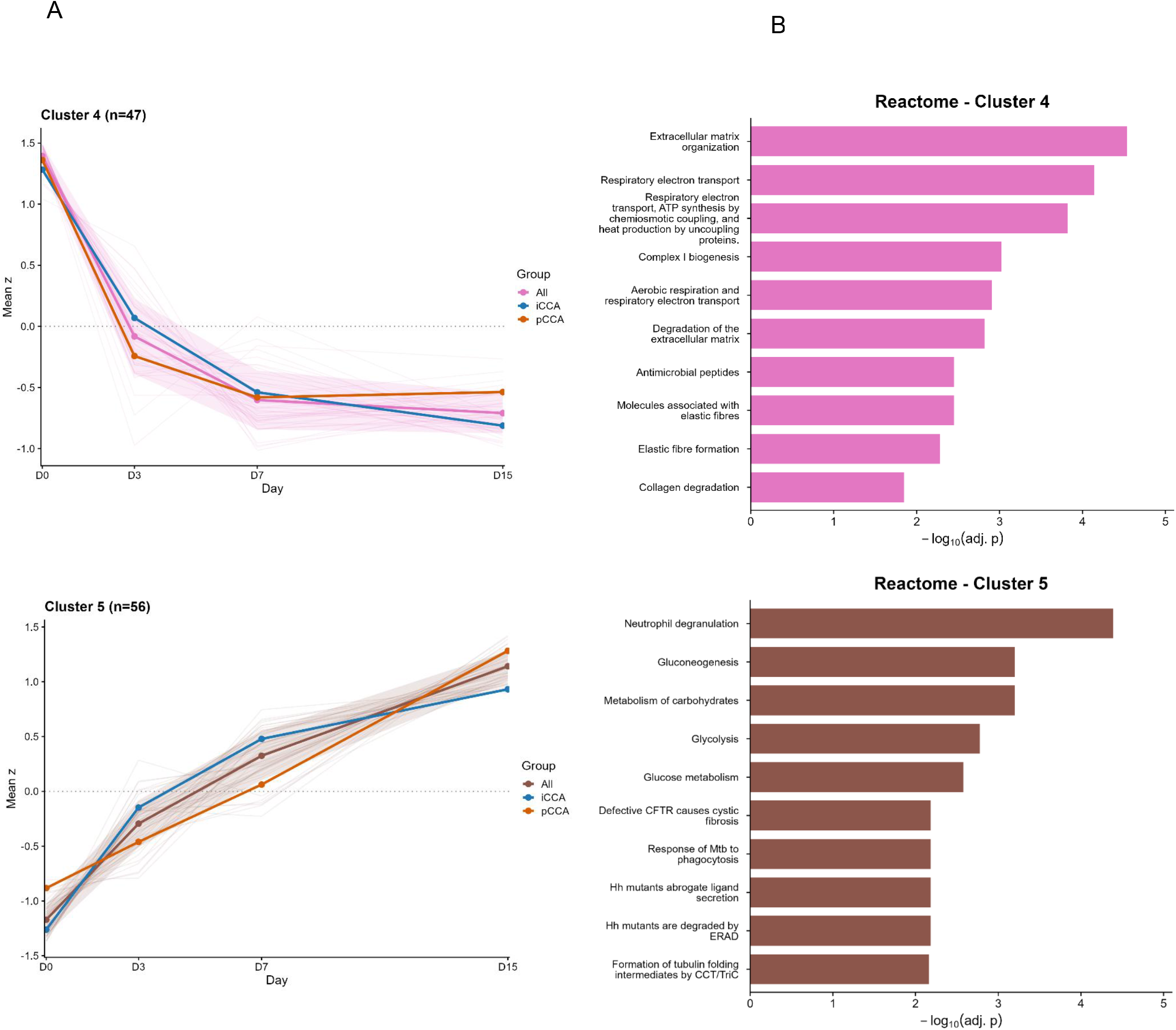
Time-varying proteins resolve into distinct temporal programmes with associated biological pathways. (A) Mfuzz soft clustering of proteins showing temporal changes during culture identified five expression patterns. Individual protein trajectories are shown as faint lines, with mean trajectories shown for all samples and by anatomical subtype; shaded areas represent ± SD. Cluster n indicates the number of proteins. (B) Reactome pathway enrichment for each temporal cluster, showing the ten most significantly enriched pathways ranked by −log10 adjusted p value.

#### 4.4.1 Early attrition of inflammatory and immune-associated proteins

Neutrophil- and plasma-associated proteins (ELANE, CTSG, S100A8/9) declined progressively from Day 3, consistent with attrition of blood- and perfusion-dependent tissue components following explantation. Closer inspection of the Reactome “neutrophil degranulation” signature, which was enriched across several Mfuzz clusters, separated classical granule and antimicrobial proteins (ELANE, CTSG, MPO, LTF, LYZ) that declined over culture from a second group of proteasomal and vesicle-trafficking proteins (PSMC3, PSMA5, RAB5B, VCP) that increased. This indicates that the Reactome term captured two biologically distinct processes rather than a single inflammatory response (Supplementary Figure 1).

#### 4.4.2 Cellular stress response

mTORC1 signalling was the most enriched pathway at every timepoint, with strong positive enrichment throughout (NES +2.07 to +2.29), driven by chaperones (HSPA5, HSPA9, HSP90B1), amino-acid transporters (SLC1A5, SLC7A5) and metabolic regulators (PHGDH, SHMT2). Cholesterol homeostasis was also positively enriched from Day 3 onwards (NES +2.05, FDR 1.69×10⁻³). Given its concurrent timing with mTORC1 activation, this is consistent with mTORC1/SREBP-driven lipid biosynthesis. Modest positive enrichment of MYC Targets V1 was also identified from Day 3 (NES +1.54, FDR 3.11 × 10⁻²). By Day 7, unfolded protein response pathways became significantly enriched (NES +1.68, FDR 2.33 × 10⁻²), alongside positive enrichment of TNFα signalling via NFκB (NES +1.96, FDR 1.31 × 10⁻²) and apoptosis (NES +1.70, FDR 2.44 × 10⁻⁴) pathways, indicating a broad inflammatory and stress-associated response emerging between Days 3 and 7. These findings were supported by temporal Mfuzz clustering, which identified a corresponding group of 65 proteins (Cluster 1) with a sustained increase in abundance, enriched for proteostasis, ER stress and mTORC1-associated pathways.

#### 4.4.3 Metabolic reprogramming

Alongside this stress response, culture drove a progressive change in cellular metabolism. Glycolysis became significantly enriched by Day 7 (NES +1.72, FDR 2.44×10⁻⁴), driven by PKM, ALDOA, PGK1, ENO1 and LDHA, and strengthened further by Day 15 (NES +2.23), accompanied by late enrichment of Hypoxia pathways (NES +1.91, FDR 6.77×10⁻³). Conversely, oxidative phosphorylation became negatively enriched from Day 7 (NES −1.55, FDR 2.44×10⁻⁴), driven by coordinated loss of respiratory-chain components (NDUFA4, NDUFV1, NDUFS2/3, COX7A2, ATP5F1C).

These findings were corroborated by temporal clustering where a 56-protein cluster (Cluster 5) containing glycolytic enzymes (PKM, GAPDH, PGK1, ENO1, PGLS, LDHA, ALDOA) progressively increased throughout culture, with Reactome enrichment confirming upregulation of glycolysis and gluconeogenesis pathways. A separate 47-protein cluster (Cluster 4) enriched for respiratory electron transport and Complex I biogenesis showed a coordinated decline in respiratory-chain proteins. A third, smaller cluster of 27 proteins (Cluster 3), enriched for mitochondrial translation, showed an initial decline followed by partial recovery by Day 15 in a subset of proteins including MRPL39, MRPL4 and MRPL44.

#### 4.4.4 Dynamic extracellular matrix remodelling

Temporal clustering identified divergent Mfuzz clusters enriched for extracellular-matrix pathways. Integration of these findings showed coordinated loss of structural stromal proteins (Cluster 4) alongside a rise in proteins governing collagen formation and cell–matrix interaction (Cluster 1). This mirrored the initial differential abundance analysis, where structural stromal proteins (FBLN1, EFEMP1, ITIH1/2) were already reduced by Day 3. Conversely, proteins involved in collagen processing and cell– matrix interaction (SERPINH1, PLOD1/3, ITGA2, ITGAV, TIMP1 and LAMC2) increased progressively during culture. These same proteins, alongside TIMP3, were also leading-edge contributors to the positive enrichment of the Hallmark epithelial– mesenchymal transition pathway within the GSEA at Day 15 (NES = 1.69, adjusted P = 5.83 × 10⁻³).

#### 4.4.5 Proteomic changes according to CCA subtype

Having established that CCA-PCTS undergo a reproducible temporal shift in their proteome, we next sought to determine whether there were any baseline differences between subtypes and if these differences persisted during culture.

When comparing subtypes directly, few individual proteins differed at baseline (Supplementary Figure 2). However, GSEA identified coordinated differences in proteomic profile, with six Hallmark programmes reaching FDR <= 0.05 (Figure 5A). Relative to pCCA, iCCA showed greater enrichment for MYC target (NES +1.99, FDR 4.1 x 10^-6^) and E2F target (NES +1.79, FDR 0.007) pathways, whereas pCCA was more enriched for the epithelial-mesenchymal transition pathway (NES -1.98, FDR 1.1 x 10^-4^). Five of the six programmes retained their direction throughout culture (Figure 5B). Oxidative phosphorylation was the exception, inverting at Day 3 before returning, consistent with the metabolic reprogramming that accompanies entry into culture (Section 4.4.3).

**Figure 5.**
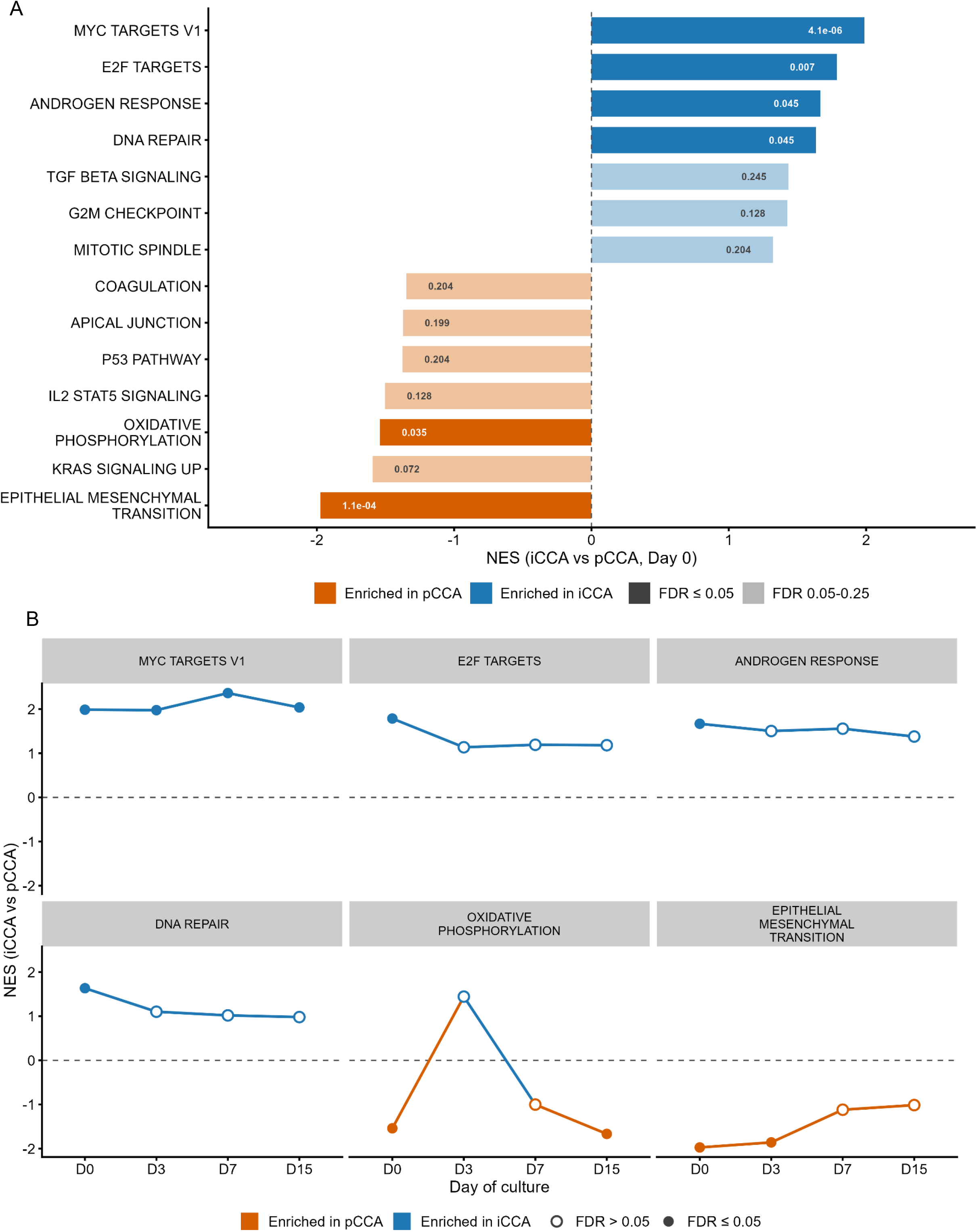
Subtype-associated proteomic programmes are retained during cholangiocarcinoma tissue slice culture. (A) Hallmark gene set enrichment analysis of the Day 0 iCCA versus pCCA proteomic comparison. Bars represent normalised enrichment scores (NES), with positive values indicating enrichment in iCCA and negative values enrichment in pCCA, and the corresponding FDR shown within each bar. All pathways reaching FDR ≤ 0.25 are displayed to show the broader enrichment landscape; solid bars indicate those meeting the formal significance threshold of FDR ≤ 0.05 and faded bars those between 0.05 and 0.25. (B) NES trajectories across Days 0, 3, 7 and 15 for the six programmes reaching FDR ≤ 0.05 at Day 0. Pathways were selected at Day 0 only and followed without re-selection at later timepoints; filled points indicate FDR ≤ 0.05 at that timepoint and open points FDR > 0.05. FDR, false discovery rate; iCCA, intrahepatic cholangiocarcinoma; NES, normalised enrichment score; pCCA, perihilar cholangiocarcinoma.

Despite these baseline differences, the overall temporal response to culture was shared. No protein showed an FDR-significant Day-by-subtype interaction (0 of 3,750), providing no evidence that the direction or magnitude of change during culture differed between subtypes. Consistent with this, Mfuzz clustering performed across the pooled cohort showed no separation of trajectories when subtype was overlaid post hoc. The proteomic response to culture was therefore common to both subtypes, while subtype-associated differences present at baseline were retained.

### 4.5 RESPONSE TO THERAPEUTIC INTERVENTION

#### 4.5.1 Response to cytotoxic treatment

Treatment with staurosporine produced a dose-dependent reduction in viability across all concentrations tested, with a corresponding increase in CC3 positivity on histology, confirming pharmacological responsiveness of PCTS to cytotoxic treatment (Figure 6a,b).

**Figure 6.**
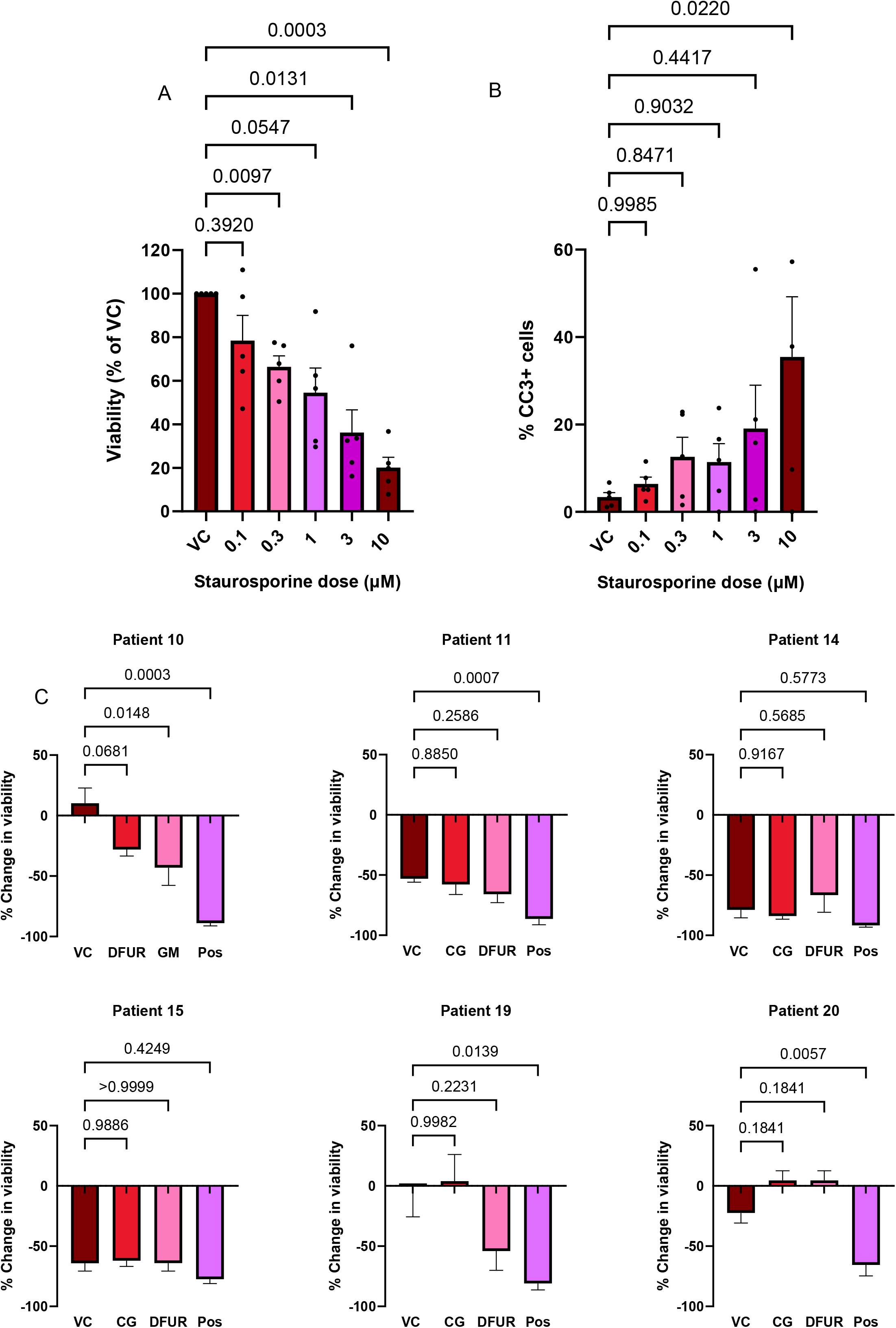
Cholangiocarcinoma tissue slices show dose-dependent cytotoxicity and heterogeneous responses to chemotherapy. (A) Tissue slice viability following 48-hour staurosporine treatment from Day 1 of culture, expressed relative to vehicle control. Data are mean ± SEM (n = 5); repeated-measures one-way ANOVA with Dunnett’s multiple-comparisons test against vehicle control. (B) CC3 positivity following staurosporine treatment. Data are mean ± SEM (n = 5); repeated-measures one-way ANOVA with Dunnett’s multiple-comparisons test against vehicle control. (C) Patient-specific responses to 5′-DFUR, gemcitabine (GM) and gemcitabine plus cisplatin (CG), expressed as percentage change relative to matched pretreatment viability. Treatment availability differed between patients. Comparisons were performed separately for each patient using one-way ANOVA with Dunnett’s multiple-comparisons test against the corresponding vehicle control. Data are mean ± SEM. CC3, cleaved caspase-3; CG, gemcitabine plus cisplatin; DFUR, 5′-deoxy-5-fluorouridine; GM, gemcitabine; Pos, staurosporine positive control; VC, vehicle control.

#### 4.5.2 Response to chemotherapy

PCTS from six patients were treated with clinically relevant CCA chemotherapy agents, including 5′-DFUR and gemcitabine ± cisplatin (40:1 ratio). Change in viability was calculated relative to matched pretreatment viability, with treatment conditions compared statistically against the corresponding vehicle control (Figure 6C). In two patients (14 and 15), high background death in vehicle-control slices precluded meaningful interpretation. Of the remaining four, one patient (Patient 10) showed a statistically significant reduction in viability following chemotherapy exposure; a further patient showed a reduction consistent with response that did not reach significance, while two showed no measurable effect, suggesting relative resistance.

## 5.0 DISCUSSION

CCA remains a formidable challenge to treat, and pre-clinical models that faithfully recapitulate the *in-vivo* tumour are critical to advancing our understanding of this disease and evaluating novel treatments. Using fresh surgical specimens, we successfully generated PCTS from 25 patients. This approach builds on prior work applying PCTS to other cancers but is among the first to comprehensively apply this methodology to CCA, including pCCA, a subtype that is technically challenging to study due to limited tissue availability and anatomical sampling constraints.

Maintaining viability of PCTS is key to maximising this model’s utility as a platform for therapeutic intervention. Our findings are consistent with other PCTS studies, including those in HCC and pancreatic ductal adenocarcinoma, where viability has been maintained for up to two weeks^13,14^. Our PCTS derived from iCCA tumours tended to show superior viability compared with pCCA and on average had a higher number of CK19+ cells. The degree of epithelial cell content is a recognised determinant of slice performance, as is the extent of tumour necrosis following neoadjuvant treatment^19^. No patients in our cohort received neoadjuvant therapy, and no other patient derived clinicopathological factors were associated with CCA PCTS viability. Beyond viability, histological analyses showed resident CD3+ and CD68+ immune cells remained identifiable throughout culture. Studies have shown resident immune cells to remain functional in PCTS for up to 48 hours^26^, whether this holds true for CCA-derived PCTS over a longer culture window remains to be established_27._

A key contribution of this study is the comprehensive proteomic characterisation over time in culture. To our knowledge, this is the first proteomic analysis of PCTS in CCA and one of the most detailed temporal datasets characterising any PCTS model to date. Previous transcriptomic studies characterising slice culture have generally been limited to the first 48–72 hours ^28–30^. Proteomic analysis offers a complementary, and arguably closer, representation of the functional tissue state, since changes at the mRNA level are not always reflected downstream at the protein level ^31,32^.

The earliest shared proteomic changes, evident from Day 3, comprised coordinated reductions in extracellular matrix and inflammatory/neutrophil-associated proteins alongside induction of stress-related chaperones, consistent with an acute response to tissue injury and transfer to an artificial culture environment. Comparable early changes have been reported in transcriptomic studies of other PCTS systems ^28^, suggesting these early changes reflect a largely generic tissue-injury response to slicing rather than a process specific to CCA.

By Day 7, the scale of proteomic change increased markedly and stabilised into a persistently altered, stress-associated state that continued through to Day 15. This stress response was dominated by mTORC1 signalling, the trigger of which cannot be attributed to a single cause. Tissue slicing and transfer to an *ex-vivo* environment represent an obvious cellular insult, and activation of growth and stress-signalling pathways in the immediate aftermath would be an expected consequence of this. However, our culture medium is also supplemented with human epidermal growth factor, an independent activator of mTORC1 and MYC signalling ^33^.

A progressive increase in glycolysis and hypoxia signatures alongside suppression of oxidative phosphorylation pathways was also seen, suggestive of a bioenergetic stress response. A small cluster of mitochondrial translation proteins showed a partial recovery by Day 15 and may represent an attempted compensatory response to progressive respiratory-chain protein loss. These changes occurred under culture conditions, predicted by mathematical modelling to achieve near-physiological internal oxygenation^34^. Comparable liver and biliary PCTS systems have nonetheless demonstrated increased HIF-1α localisation under similar culture conditions^14^. This suggests that even optimised culture conditions may not fully abolish diffusion-limited hypoxic stress, particularly in densely desmoplastic tumour types such as CCA. The use of dynamic flow systems may reduce tissue hypoxia relative to static culture ^35,36^, but add considerable technical complexity and are often bespoke to individual laboratories, limiting their scalability and reproducibility.

We also observed divergent trajectories among ECM-associated proteins. Structural stromal proteins declined progressively from Day 3, while proteins governing collagen processing and cell–matrix interaction increased. This pattern parallels transcriptomic findings in other PCTS systems^28^, as well as reports of spontaneous fibrogenesis during prolonged culture of human liver slices^37^. Notably, this ECM remodelling occurred despite preservation of overall tissue architecture on histology, implying that molecular-level change may precede any visible structural deterioration, or that active-matrix remodelling helps maintain tissue architecture *ex-vivo*.

Despite few individual proteins differing between anatomical subtypes at baseline, pathway-level analysis identified coordinated subtype-associated proteomic programmes, including greater MYC- and E2F-associated enrichment in iCCA and greater epithelial–mesenchymal transition enrichment in pCCA. Five of six programmes retained their direction during culture, while no protein showed an FDR-significant Day-by-subtype interaction. Together, these findings suggest that hPCTS retain subtype-associated proteomic organisation alongside a shared response to ex vivo culture. However, the subtype analysis included only five donors per group, limiting power to detect interaction effects, and bulk proteomics cannot distinguish intrinsic tumour-cell differences from variation in epithelial, stromal or immune composition.

PCTS are an attractive model for evaluating drug responses^38^. We confirmed pharmacological responsiveness using STS, which produced a dose-dependent reduction in viability and served as a robust positive control. Responses to clinically relevant chemotherapy agents were more variable. In two cases, high background death in VC slices precluded meaningful comparisons, a potential issue in PCTS-based therapeutic studies where baseline viability and tissue quality can dominate assay interpretability ^24^. Among evaluable cases, only one patient showed a statistically significant reduction in viability with chemotherapy treatment, while others showed partial or no measurable effect. These mixed responses may reflect true biological resistance but could reflect limited drug penetration/metabolism or simply insufficient dynamic range in some slices. Notably, however, the patient with the greatest ex-vivo chemosensitivity was also the only individual alive without confirmed recurrence at last follow-up. This single-case observation should be interpreted cautiously, given the influence of multiple clinical variables on outcome, but is consistent with the broader premise that PCTS can capture clinically relevant differences in drug sensitivity ^21,39^.

Our study has several limitations. CCA is rare, and despite extended accrual, sample numbers remain modest. The inherently desmoplastic and spatially heterogeneous nature of CCA introduces variability in slice composition and viability that can limit throughput and assay reproducibility. We acknowledge some of these challenges may be CCA-specific and not generalisable to all tumour types.

While our proteomic data demonstrate a clear temporal response to culture, bulk tissue proteomics cannot resolve which cellular compartment underpins the proteomic changes described, and direct measurement of oxygen tension within the slice was not performed, meaning the conclusions drawn on hypoxic stress signatures remain inferential. A contrast with a non-tumour dataset would also help establish whether the changes observed are cancer-specific or represent a general feature of all PCTS culture. Furthermore, the functional consequences of these proteomic changes on drug sensitivity and treatment response remain unclear. The metabolic shift away from oxidative phosphorylation and towards glycolysis, together with sustained proteostatic stress signalling, could plausibly alter drug sensitivity independent of a tumour’s native biology. Equally, the extracellular matrix remodelling described above may influence drug penetration through the slice. Distinguishing genuine tumour drug sensitivity from culture-stage-dependent artefacts will require drug dosing to be performed and compared across multiple culture timepoints, alongside targeted assessment of drug-transporter and metabolising enzyme expression.

Finally, whilst we show resident immune cells remain identifiable in culture, further work is required to assess their functionality. A general limitation of PCTS culture is the lack of any systemic vascular inputs, including circulating immune cells, making studies of immune cell recruitment and TME modulation challenging, although co-culture with patient-matched PBMCs offers a potential solution^40,41^.

In summary, we establish CCA PCTS as a viable *ex-vivo* platform capable of maintaining tumour architecture and subtype identity for up to 15 days. We show that PCTS undergo a shift in proteomic profile characterised by stress responses and metabolic rewiring that stabilises by day 7. We also demonstrate pharmacological responsiveness of PCTS to cytotoxic treatment with variability in chemotherapy response. Collectively, these findings support further development of PCTS for mechanistic and therapeutic studies in biliary tract cancer.

## ABBREVIATIONS

3Rs: Replacement, Reduction and Refinement (of animals in research)
CC3: Cleaved caspase-3
CCA: Cholangiocarcinoma
DAB: Diaminobenzidine
DDA: Data-dependent acquisition
DFUR: 5′-deoxy-5-fluorouridine
DIA-NN: Data-independent acquisition by neural networks
DMSO: Dimethyl sulfoxide
FOV: Field of view
GemCis: Gemcitabine plus cisplatin
GSEA: Gene set enrichment analysis
HCC: Hepatocellular carcinoma
hEGF: Human epidermal growth factor
HRP: Horseradish peroxidase
HTA: Human Tissue Authority
iCCA: Intrahepatic cholangiocarcinoma
LC–MS: Liquid chromatography–mass spectrometry
MSigDB: Molecular Signatures Database
MTS: Colorimetric cell viability assay (CellTiter 96® AQueous One Solution Cell Proliferation Assay)
NES: Normalised enrichment score
PBMC: Peripheral blood mononuclear cells
pCCA: Perihilar cholangiocarcinoma
PCTS: Precision-cut tissue slices
REC: Research Ethics Committee
SCX: Strong cation exchange
SP3: Single-pot solid-phase-enhanced sample preparation
SREBP: Sterol regulatory element-binding protein
STS: Staurosporine
SWATH-MS/SWATH-DIA: Sequential window acquisition of all theoretical fragment-ion mass spectra (data-independent acquisition)
TME: Tumour microenvironment
UW: University of Wisconsin
WEM: Williams’ E Media

## ACKNOWLEDGEMENTS

We thank the patients who consented to donate tissue, without whom this research would not have been possible.

## Author contributions: Timothy M Gilbert

Conceptualization, Methodology, Investigation, Formal analysis, Data curation, Writing – Original draft preparation, Writing – Review & editing. **Owen McGreevy**: Methodology, Investigation, Formal analysis, Data curation, Writing – Original draft preparation, Writing – Review & editing. **Maria-Danae Jessel**: Investigation, Formal analysis. **Roz Jenkins**: Investigation (proteomics). **Mohamed Bosakhar**: Investigation. **Marc Quinn, Lawrence O’Leary**: Resources. **Anthony Evans**: Formal analysis (bioinformatics). **Timothy Andrews**: Investigation (histopathology). **Rafael Diaz-Nieto, Robert P Jones, Stephen Fenwick**: Resources. **William Greenhalf, Daniel Palmer**: Supervision, Writing – Review & editing. **Hassan Z Malik**: Resources, Supervision, Writing – Review & editing. **Christopher Goldring**: Conceptualization, Supervision, Funding acquisition, Writing – Review & editing. **Laura Randle**: Conceptualization, Methodology, Investigation, Supervision, Funding acquisition, Writing – Review & editing. All authors reviewed and approved the final version of the manuscript for submission.

