## Supplementary Material for "A PATIENT-DERIVED EX VIVO TISSUE MODEL OF CHOLANGIOCARCINOMA USING PRECISION-CUT TISSUE SLICES"

### ***METHODS AND MATERIALS***

#### *TISSUE COLLECTION*

Patients were identified through the specialist hepatobiliary multidisciplinary team (MDT) at Liverpool University Hospitals NHS Foundation Trust and were approached pre-operatively for consent. Tissue donation was conducted in accordance with University of Liverpool and HTA policies. Tissue was collected from the operating theatre following removal of the surgical specimen; a bench-side dissection of the resected liver was performed in theatre and a section of tumour tissue was removed for research. Co-ordination with the histopathology team was undertaken to ensure sufficient specimen integrity was retained for assessment of the patient resection margin.

#### *GENERATION OF CCA hPCTS*

Obtained specimens were transferred from theatre to the Human Liver Research Facility (University of Liverpool). Tumour sections were removed from University of Wisconsin buffer using sterile forceps and placed on a sterile petri dish with sufficient buffer to keep the specimen moist. Samples were processed using a 5 mm diameter coring drill (Alabama Research and Development, Munford, AL, USA) to produce multiple cylindrical cores, which were inserted into a Krumdieck Tissue Slicer at 4 °C to generate multiple 250 µm slices (arm speed 4, blade speed 4). Slices generated from multiple cores were collected from the glass trap and transferred into supplemented WEM for 1 h at 37 °C to allow recovery. Slices from multiple cores were pooled in the reservoir solution and allocated randomly to culture conditions, giving a random sample across the tumour rather than from any single core.

#### *EX-VIVO CULTURE OF CCA hPCTS*

Following 1 h of initial recovery, the required number of hPCTS were transferred to fresh 24-well plates placed on 12 mm, 0.4 µm pore size Millicell® Cell Culture Inserts (Merck, Cat# PICM01250), placed on an orbital shaker (SLS Lab Basics Digital Orbital Shaker, Cat# SLS6060) at 95 rpm and incubated in normoxic conditions at 37 °C and 5% $CO_2$. The use of porous organotypic inserts lifts the tissue slices off the bottom of the culture plate, holding the tissue slice at a fixed depth within the culture well and helps to maintain a consistent air-liquid interface. Each culture well contained 450 µL of fresh supplemented WEM with 200 ng/mL hEGF (Corning, Cat# 354052).

#### *MTS VIABILITY ASSAY*

Individual MTS assays were performed by removing selected tissue slices from culture and placing them in fresh 24-well culture plates (no insert) containing 400 µL of supplemented WEM (without hEGF) plus 80 µL of CellTiter 96® AQueous One Solution reagent (Promega, Cat# G3582). Wells containing reagent but no tissue served as blanks. Plates were incubated at 37 °C and 5% $CO_2$, protected from light, and agitated on an orbital shaker. After 3 h, 200 µL of the MTS/WEM solution from each well was transferred to a corresponding well of a 96-well plate and absorbance read at 490 nm (Varioskan Flash 2.4.5; Thermo Fisher Scientific, Cat# VLBL00GD2). Mean blank absorbance was subtracted from each sample value to remove background. Following the assay, slices from the timepoint of interest were washed and returned to culture, snap-frozen, or fixed in 10% neutral-buffered formalin (Sigma-Aldrich, Cat# HT501128) as required for downstream analysis.

#### *hPCTS THERAPEUTIC DOSING*

CCA hPCTS were prepared as described and maintained overnight in WEM for recovery. On Day 1, slices were transferred to fresh WEM and treated for a minimum of 48 h with either STS, GemCis combination therapy, or DFUR, with a complete medium exchange at 24 h. STS was prepared from a 2 mM stock in 100% DMSO and diluted to 0.1, 0.3, 1, 3 and 10 µM in WEM, with 50% DMSO as a positive control. Gemcitabine was prepared from a 20 mM stock in 100% DMSO and cisplatin from a 2.5 mM stock in 0.9% saline. For GemCis combination treatments, each gemcitabine concentration (1, 3, 10, 30, 100 μM) was co-administered with the corresponding cisplatin concentration (0.025, 0.079, 0.25, 0.79, 2.5 μM) to achieve a half-log incremental dosing series for both agents, ensuring total DMSO did not exceed 0.5%. DFUR was prepared from a 20 mM stock in 100% DMSO and diluted to 1, 3, 10, 30 and 100 μM in WEM, with 10 μM STS as a positive control. Vehicle controls (0.5% DMSO and saline) and untreated WEM controls were included on each plate. STS-, GemCis- and DFUR-treated slices were allocated to separate 24-well plates, with three replicates per condition. After the 48 h incubation, hPCTS were collected for downstream analyses of therapeutic efficacy and tissue viability.

### *FFPE BLOCK PREPARATION, MICROTOMY & SECTIONING*

#### *FFPE processing and sectioning*

Slices fixed in 10% neutral-buffered formalin (Sigma-Aldrich, Cat# HT501128) were dehydrated through 70%, 90% and 100% ethanol, cleared in 100% xylene, and embedded flat in paraffin wax for transverse sectioning. Blocks were chilled tissue-side down on ice for 10 min, and 4 μm sections cut on a microtome (Leica Biosystems, Reichert-Jung Biocut 2030), floated on a 37 °C water bath, mounted on Superfrost™ Plus slides (Fisher Scientific, Cat# 10149870) and dried overnight at 40 °C.

#### *H&E staining*

Sections were deparaffinised in xylene (5 min), taken through graded ethanol (100%, 95%, 70%), stained in haematoxylin (Sigma-Aldrich, Cat# H9627; 10 min), differentiated in 0.25% acid alcohol and blued in Scott's tap water, counterstained in eosin (Sigma-Aldrich, Cat# 318906; 2 min), dehydrated, cleared in xylene and mounted in DPX (Sigma-Aldrich, Cat# 06522).

#### *Immunohistochemistry*

Deparaffinisation and heat-induced antigen retrieval were performed on a Dako PT-Link using low- or high-pH Target Retrieval Buffer (Agilent, Cat# S236984-2 or S236784-2) according to antibody requirements. Unless otherwise stated, reagents were from the Dako REAL EnVision Detection System Peroxidase/DAB+, Rabbit/Mouse-HRP kit (Agilent, Cat# K500711-2). Sections were washed in TBS-T, blocked in peroxidase blocking solution (10 min) and incubated with primary antibody: CC3 (1:300, 2 h RT), Ki67 (1:1000, 1 h RT), CK19 (1:400, 1 h RT), CD3 (1:200, 1 h RT) or CD68 (1:200, 1 h RT). HRP-conjugated secondary antibody was applied (1 h RT), signal developed with DAB (10 min), and sections counterstained with haematoxylin, dehydrated, cleared and mounted in DPX. All slides for a given marker were stained within a single batch and processed at the same time.

*Digital pathology*

Cover-slipped slides were reviewed for tissue and staining quality and scanned on an Aperio CS2 ScanScope at 20× to generate whole-slide brightfield (H-DAB) CZI file images and analysed in QuPath v0.6.0. A single set of colour deconvolution stain vectors was estimated from representative tissue and applied consistently across all slides. Each slide was analysed using the Positive Cell Detection function, with single-intensity thresholds set with reference to matched batch-level positive and negative control tissue (CC3 0.2, CK19 0.18, Ki67 0.22, CD3 0.1, CD68 0.25) and object feature smoothing applied over a 25 μm radius. Percentage positivity was calculated for each slice as the number of positive cell detections divided by the total number of cell detections.

### *PROTEOMIC SAMPLE PREPARATION AND SWATH-MS ACQUISITION*

#### *Spectral library preparation*

A project-specific spectral library was generated from pooled bulk resection tissue protein from nine CCA patients (iCCA and pCCA), five of whom also contributed hPCTS to the SWATH-DIA sample set. Pooled protein (1.5 mg) in urea lysis buffer was processed in solution without bead capture, as a higher protein input was required than the bead-based route allowed. Protein was reduced with 5 mM dithiothreitol (DTT; 30 min, 60 °C, 1000 rpm), cooled, alkylated with 20 mM iodoacetamide (30 min, dark, room temperature), and quenched with 5 mM DTT (15 min). Digestion was performed in two stages. Trypsin/Lys-C (75 µg) was added in 6 M urea and incubated for 3 h at 37 °C (1000 rpm), after which the urea was diluted fivefold with water to 1 M and digestion continued overnight at 37 °C. Lys-C enables cleavage under high-urea denaturing conditions and, with subsequent trypsin activation, increases cleavage efficiency at lysine residues. The peptide mixture was prefractionated by strong cation exchange (SCX) chromatography (200 × 4.6 mm, 5 µm, 300 Å; Poly LC, Columbia, MD). Peptides were adjusted to 5 mL with SCX buffer A, acidified to pH 3 with phosphoric acid, and clarified (17,000 × g, 3 min, room temperature) before loading. Eighty 1 mL fractions were collected, of which approximately 40 contained peptides; these were dried by vacuum centrifugation (SpeedVac, Eppendorf UK Ltd, Stevenage, UK), resuspended in 1 mL 0.1% trifluoroacetic acid (TFA), clarified again, desalted on an Agilent mRP-C18 column, and each analysed individually by DDA over a 150 min gradient.

*CCA tumour PCTS sample preparation (SWATH-DIA)*

Tumour hPCTS were stored at −80 °C. For each sample, approximately 10 mg tissue (equivalent to approximately two slices) was homogenised by hand with a sterile single-use polypropylene pestle in 130 µL ice-cold lysis buffer (6 M urea, 1 M ammonium bicarbonate, 0.5% sodium deoxycholate) with protease inhibitors (Thermo Scientific, Cat# 78430). Lysates were clarified at 17,000 × g for 15 min at 4 °C and the supernatant retained. Protein was quantified by Bradford assay at $A_{595}$ against a bovine serum albumin (BSA) standard curve, with samples diluted 1:5 to 1:40 to fall within the linear range of the assay. For each sample, 100 µg total protein was made up to 100 µL in 100 mM ammonium bicarbonate, reduced with 5 mM DTT (30 min, 60 °C), cooled, alkylated with 20 mM iodoacetamide (30 min, dark, room temperature), and quenched with a further 5 mM DTT (15 min). Proteins were captured by SP3 using a 1:1 mix of hydrophilic and hydrophobic carboxylate-modified Sera-Mag SpeedBeads (Cytiva) at 70% ethanol (v/v) and mixed for 1 h at 24 °C. Beads were washed three times with 80% ethanol and, for on-bead digestion, resuspended in 100 mM ammonium bicarbonate containing 2.4 µg sequencing-grade trypsin and incubated overnight at 37 °C. Peptides were recovered magnetically, acidified to pH 2–3 with TFA, desalted by on-line injection onto an Agilent mRP-C18 column, dried, and reconstituted in 0.1% formic acid for LC-MS/MS.

*LC–MS and data acquisition (library fractions and SWATH samples)*

Peptides (0.5–1 µg) were loaded onto a Symmetry C18 nanoAcquity trap column (Waters) and separated on a bioZen XB-C18 column (75 µm × 250 mm; Phenomenex) over a linear gradient of 2–50% acetonitrile in 0.1% formic acid at 300 nL/min, on a TripleTOF 6600 (SCIEX). Spectral library fractions were separated over a 150 min gradient and acquired by information-dependent acquisition (IDA; 400–1500 m/z, top-25 precursors per 2.8 s cycle, 20 s dynamic exclusion), and searched in ProteinPilot 5.1 (Paragon algorithm, SCIEX) against UniProt Swiss-Prot (human, 82,492 entries), retaining proteotypic peptides with carbamidomethylated cysteines at 1% FDR against a reversed decoy database. The resulting spectral library was exported using PeakView. Individual hPCTS samples were separated over a 120 min gradient in two experimental batches and acquired in SWATH mode using 100 variable windows across a precursor range of 400–1500 m/z, with fragment ions recorded across 100–1650 m/z (3.1 s cycle).

*DIA-NN processing and quantification*

MS data files were converted to DIA-NN format prior to processing. The PeakView-exported spectral library and the UniProt Swiss-Prot human FASTA (as used for the ProteinPilot search) were loaded, with re-annotation enabled. Digestion was set to Trypsin/P with one missed cleavage. Carbamidomethylation of cysteine and N-terminal methionine excision were applied as fixed modifications, with no variable modifications, to reflect the basal peptide form. Precursor and fragment m/z ranges were set to 400–1500 and 100–1650 respectively, with peptide and charge states left at default. Library generation used smart profiling and the neural network classifier was run in double-pass mode. Cross-run normalisation was RT-dependent (default), the quantification strategy was left at default, and match-between-runs, use of isotopologues, and unrelated-run calibration were enabled. In double-pass mode, the spectral library is first matched to the DIA data, after which the library is rebuilt from the raw data to recover lower-abundance proteins identified with sufficient confidence, yielding higher-confidence identifications than single-pass processing. Protein identifications were controlled at 1% FDR and the protein-group (PG) matrix exported for downstream analysis.

*BIOINFORMATIC & STATISTICAL ANALYSIS*

*CLINICAL AND ROUTINE LABORATORY STATISTICAL ANALYSIS*

Analyses were performed in SPSS Statistics v22 (IBM), GraphPad Prism v10.4 (GraphPad Software) and R v4.4.2. Normality was assessed with the Shapiro-Wilk test. Normally distributed variables are reported as mean ± SD and compared with two-sided Student's t-tests; non-normal data are reported as median (interquartile range) and analysed with non-parametric tests. Metabolic activity was assessed by MTS assay at Days 0, 3, 7, 11 and 15 of culture; missing timepoints reflected tissue availability rather than slice failure. Raw MTS absorbance was modelled by restricted maximum likelihood with culture day as a fixed effect and patient as a random effect, with the Geisser-Greenhouse correction applied and each timepoint compared with Day 0 by Dunnett's test. Model residuals were assessed graphically. Immunohistochemical marker positivity (CC3, Ki67, CK19) was quantified at Days 0, 3, 7 and 15 as repeated measures within patient. CK19 and CC3, which were non-normally distributed with complete data, were analysed by the Friedman test with Dunn's multiple comparisons against Day 0; Ki67, which contained missing timepoints, was analysed by a mixed-effects model with Dunnett's comparisons against Day 0. Baseline (Day 0) MTS absorbance was compared between iCCA and pCCA by Welch's two-sided t-test

(Supplementary Table 4). Significance was set at $p \leq 0.05$ unless otherwise stated. Therapeutic-response experiments were analysed using one-way analysis of variance with Dunnett's multiple-comparisons test, with each treatment condition compared against the corresponding vehicle control. Staurosporine dose-response data were analysed across patients as repeated measures. Chemotherapy responses were analysed separately for each patient, with treatment conditions compared with the matched vehicle control. Data are presented as mean ± SEM.

*MS data QC and imputation*

The DIA-NN protein-group intensity matrix was imported into R v4.4.2 and log2-transformed. Quality control comprised missingness and value-distribution heatmaps annotated by donor, day (D0, D3, D7, D15), MS batch (two batches) and subtype (iCCA, pCCA). Missing values were imputed by random forest (imp4p::impute.RF) with CCA subtype supplied as the conditioning variable, so that imputation was informed by subtype structure. Protein annotations (Protein.Group, gene symbol, description) were merged to generate the final assay matrix used for downstream testing and visualisation.

*Filtering of the Day 0 subtype comparison*

Because imputation was conditioned on CCA subtype, imputed values carry subtype information and can contribute to an apparent difference in a direct subtype comparison. This does not introduce the same circularity for temporal contrasts, where the conditioning variable differs from the variable being tested. The Day 0 subtype comparison was therefore restricted, before model fitting, to protein groups with observed pre-imputation quantification in at least three of five samples within each subtype (≥60% detection per subtype), retaining 3,750 of 4,578 protein groups.

*Differential abundance*

Differential abundance was modelled in limma on the log2 protein-group matrix, with repeated measures across donors accounted for by duplicateCorrelation (block = Donor) passed to lmFit, followed by empirical Bayes moderation (eBayes) with trend and robust options. For the overall analysis an additive design (Batch + Subtype + Day) was used, and all pairwise contrasts among Days 0, 3, 7 and 15 were tested, with proteins considered differentially abundant at Benjamini–Hochberg FDR ≤ 0.05 and |log2 fold-change| ≥ 0.5.

Subtype comparisons used an interaction design (Batch + Day x Subtype) with treatment contrasts, Day 0 as the reference day and perihilar as the reference subtype, fitted on the filtered protein set described above with the same donor blocking and moderation. The subtype coefficient at the reference day gives the Day 0 iCCA versus pCCA contrast, and the sum of that coefficient and the corresponding interaction term gives the subtype difference at each later day. Whether the temporal response to culture differed by subtype was assessed by a joint moderated F test across the three Day-by-subtype interaction coefficients, with Benjamini-Hochberg control.

Principal component analysis (prcomp) was performed on the imputed log2 matrix for the overall cohort and within subtypes; batch-adjusted score plots (removeBatchEffect) were used for display only and not for hypothesis testing. Volcano plots and heatmaps of variable proteins and significant differentially abundant proteins (DAPs) were generated in R.

*Gene set enrichment analysis*

GSEA was performed with fgsea using the MSigDB Hallmark collection retrieved via msigdbr (Homo sapiens). For each contrast, limma results were collapsed to one row per gene symbol by selecting the entry with the maximum absolute moderated t-statistic, and the signed moderated t provided the ranking metric. Enrichment was computed against the set of quantified genes with a minimum set size of 10 and a maximum of 500, using 10,000 permutations and a fixed random seed so that reported values are reproducible, with significance at FDR <= 0.05 (Benjamini-Hochberg). Outputs comprised normalised enrichment scores (NES), adjusted p-values and leading-edge members, visualised as NES heatmaps across contrasts, running-enrichment curves and NES trajectories across days.

The workflow was run for the overall temporal analysis and the subtype comparison. For the subtype analysis, enrichment was computed on the filtered Day 0 contrast. Figure 5A displays pathways with FDR ≤ 0.25 to show the broader enrichment landscape, while pathways meeting the formal significance threshold of FDR ≤ 0.05 at Day 0 were carried forward and their NES tracked across Days 3, 7 and 15 without re-selection.

*Temporal proteomic analysis*

Temporal behaviour was modelled with Mfuzz (v2.64.0) on the overall cohort. The input comprised the union of proteins significant in any of the Day 3, 7 or 15 versus Day 0 limma contrasts (FDR ≤ 0.05) with an additional effect-size filter (|log2FC| ≥ 0.5), giving 232 proteins. Log2 intensities were z-standardised across D0, D3, D7 and D15 using day means. The fuzzifier m was estimated with mestimate and the number of clusters set to K = 5, guided by the elbow criterion. Clustering used fuzzy c-means, with core members defined at membership ≥ 0.7, and trajectories plotted with proportional spacing of days. Clustering was performed once on the overall cohort; mean cluster trajectories were then displayed for the whole cohort and, as overlays, separately for iCCA and pCCA to show subtype-specific behaviour within the shared clusters. Cluster members were mapped to Entrez IDs (org.Hs.eg.db) and tested for pathway enrichment with ReactomePA (Benjamini-Hochberg FDR control), with the top enriched Reactome pathways reported per cluster.

*Software*

Analyses were performed in R v4.4.2 (Bioconductor 3.19). Missing protein-group intensities were imputed with imp4p v1.2 (random forest, based on the missForest algorithm), and differential abundance was modelled with limma v3.60.6. Gene set enrichment analysis used fgsea v1.30.0 with Hallmark gene sets from msigdbr v26.1.0, and Reactome enrichment used ReactomePA v1.48.0 and clusterProfiler v4.12.6 with org.Hs.eg.db v3.19.1 and AnnotationDbi v1.66.0. Temporal clustering used Mfuzz v2.64.0 (Biobase v2.64.0). Figures were generated with ggplot2 v4.0.3, pheatmap v1.0.12, cowplot v1.2.0, ggrepel v0.9.6, RColorBrewer v1.1.3, scales v1.4.0 and magick v2.9.1, with data handling in data.table v1.18.4. Clinical statistics used SPSS v22 and GraphPad Prism v10.4.

**Supplementary Tables and Figures**

**Supplementary Table 1.** Composition of 10× Krebs–Henseleit buffer stock and 1× working Krebs–Henseleit buffer solutions.

| **10x KHB** | | | |
|---|---|---|---|
| **Chemical** | **Molecular weight (g/mol)** | **Amount required / 5L Total** | **Final concentration in 10× stock** |
| **$dH_2O$** | | 2L | |
| **$CaCl_2·2H_2O$** | 147.0 | 18.35 g | 25 mM |
| **KCl** | 74.55 | 18.65 g | 50 mM |
| **NaCl** | 58.44 | 345 g | 1180 mM |
| **$MgSO_4·7H_2O$** | 246.48 | 13.55 g | 11 mM |
| **$KH_2PO_4$** | 136.09 | 8.15 g | 12 mM |
| 10× KHB was prepared as two separate solutions, with $CaCl_2·2H_2O$ dissolved separately from the remaining salts before combining and making up to a final volume of 5 L with $dH_2O$. | | | |

| 1X KHB | | |
|---|---|---|
| **Solution** | **Volume** | **Stock Concentration** |
| **KHB** | 100 ml | 10X |
| **Sodium bicarbonate** | 25 ml | 1 M |
| **Glucose** | 25 ml | 1 M |
| **HEPES** | 10 ml | 1 M |
| **$dH_2O$** | 840 ml | |

**Supplementary Table 2.** Composition of supplemented Williams' E medium.

*Sterile-filtered components**

| Supplemented Williams' E media (WEM) | | | |
|---|---|---|---|
| **Solution** | **Volume** | **Final concentration** | **Stock concentration** |
| **Nicotinamide** | 6 mL | 12 mM | 100 mM * |
| **L-Ascorbic Acid** | 6 mL | 175 µM | 14.5 mM * |
| **Sodium bicarbonate** | 13.4 mL | 0.225% w/v<br>(26.8 mM) | 7.5%<br>1 M * |
| **Glucose** | 13.9 mL | 27.37 mM | 1 M * |
| **HEPES (Sigma-Aldrich, Cat# H0887-100ML)** | 10 mL | 20 mM | 1 M |
| **Sodium pyruvate (Gibco, Cat# 11360070)** | 5 mL | 1 mM | 100 mM |
| **L-Glutamine (Gibco, Cat# 25030081)** | 5 mL | 2 mM | 200 mM |
| **Penicillin-Streptomycin (Gibco, Cat# 15140-122)** | 2 mL | 0.4% | |
| **ITS + Premix (Corning, Cat# 354352)** | 5 mL | 1% | 100% |
| **Williams E Media** | 433.7ml | - | - |

| *Patient No.* | Experiments | | | | Clinical characteristics | | | | | | | | | | Operation | Histology | | | | | | | |
|---|---|---|---|---|---|---|---|---|---|---|---|---|---|---|---|---|---|---|---|---|---|---|---|
| | Ti | P (lib) | P (ti) | Th | Age | Sex | BMI | Smoker | Diabetes | ETOH xs | PSC | Pre-op Jaundice | Biliary drainage | Pre-op PVE | Operation | Type | B-C score | Tumour size (mm) | Grade | T stage | N stage | LVI | PNI |
| *Patient 1* | Ti | | | | 44 | Female | 33.4 | No | No | No | No | No | No | No | Left Hepatectomy, caudate lobectomy and radical bile duct resection | pCCA | 3b | 65x48 | Poor | 2b | 1 | Yes | Yes |
| *Patient 2* | Ti | P (lib) | | | 75 | Male | 29.6 | No | No | No | No | Yes | ERCP | No | Left Hepatectomy, caudate lobectomy and radical bile duct resection | pCCA | 3b | 24x19 | Poor | 2b | 0 | Yes | Yes |
| *Patient 3* | Ti | P (lib) | | | 68 | Female | 23.8 | No | Yes | No | No | No | No | No | Left Hepatectomy, caudate lobectomy and radical bile duct resection | pCCA | 3b | 43x35 | Poor | 2b | 0 | Yes | Yes |
| *Patient 4* | Ti | P (lib) | P (ti) | | 78 | Male | 26.9 | Yes | No | No | No | No | No | No | Left Hepatectomy | iCCA | n/a | 26x26 | Moderate | 1a | x | No | Yes |
| *Patient 5* | Ti | P (lib) | P (ti) | | 64 | Female | 35.2 | Ex | Yes | No | No | No | No | No | Right posterior sectionectomy | iCCA | n/a | 45x31 | Moderate | 2 | x | No | No |
| *Patient 6* | Ti | P (lib) | P (ti) | | 75 | Male | 25.2 | No | Yes | Yes | No | Yes | ERCP | No | Right hepatectomy and radical bile duct resection | pCCA | 3a | 25x15 | Poor | 2b | 0 | Yes | Yes |
| *Patient 7* | Ti | P (lib) | P (ti) | | 62 | Male | 20.5 | No | No | No | No | Yes | ERCP | No | Ext Left Hepatectomy, caudate lobectomy and radical bile duct resection | pCCA | 3b | 20x24 | Well | 2a | 0 | Yes | Yes |
| *Patient 8* | Ti | P (lib) | P (ti) | | 74 | Female | 26.8 | No | No | No | No | No | No | No | Segment V resection | iCCA | n/a | 30x44 | Poor | 1a | x | No | No |
| *Patient 9* | Ti | | P (ti) | | 74 | Male | 27.2 | No | No | No | No | Yes | ERCP | No | Radical Bile duct resection | pCCA | 2 | 15x18 | Poor | 1 | 0 | No | No |
| *Patient 10* | Ti | P (lib) | | Th | 38 | Female | 28.1 | No | No | No | No | No | No | No | Left Hepatectomy | iCCA | n/a | 46x45 | Moderate | 1a | x | No | No |
| *Patient 11* | Ti | | | Th | 75 | Male | 25.2 | No | No | No | No | No | No | No | Left Hepatectomy | iCCA | n/a | 72x65 | Poor | 3 | x | Yes | No |
| *Patient 12* | Ti | P (lib) | | | 80 | Male | 27.4 | No | No | No | No | No | No | No | Left Hepatectomy, caudate lobectomy and radical bile duct resection | pCCA | n/a | 85x50 | Moderate | 2 | x | No | Yes |
| *Patient 13* | | | | | 75 | Male | 22.1 | No | No | No | No | No | No | No | Segment IV resection | iCCA | n/a | 32x27 | Moderate | 1a | x | No | No |
| *Patient 14* | | | | Th | 80 | Female | 31.8 | No | No | No | No | No | No | No | Left Hepatectomy | iCCA | n/a | 36x53 | Moderate | 1a | x | No | No |
| *Patient 15* | | | | Th | 61 | Male | 31.9 | No | No | No | No | Yes | ERCP | No | Left Hepatectomy, caudate lobectomy and radical bile duct resection | pCCA | 3b | 57x32 | Poor | 2b | 1 | No | Yes |
| *Patient 16* | Ti | | P (ti) | | 70 | Female | 22.3 | No | No | No | No | Yes | ERCP | No | Left Hepatectomy, caudate lobectomy and radical bile duct resection | pCCA | 3b | 18x16 | Poor | 2b | 1 | Yes | Yes |
| *Patient 17* | Ti | | P (ti) | | 59 | Female | 29.6 | No | No | No | No | Yes | ERCP | No | Right hepatectomy and radical bile duct resection | pCCA | 3a | 49x35 | Poor | 3 | 1 | Yes | Yes |
| *Patient 18* | Ti | | | | 72 | Male | 23.3 | No | No | No | No | Yes | ERCP | No | Left Hepatectomy, caudate lobectomy and radical bile duct resection | pCCA | 3b | 24x30 | Moderate | 2b | 0 | Yes | Yes |
| *Patient 19* | Ti | | P (ti) | Th | 81 | Female | 20.5 | No | No | No | No | No | No | No | Right hepatectomy | iCCA | n/a | 140x115 | Moderate | 2b | x | Yes | No |
| *Patient 20* | | | | Th | 70 | Male | 34 | No | Yes | Yes | No | No | No | No | Segment VII Resection | iCCA | n/a | 72x60 | Moderate | 1b | x | No | No |
| *Patient 21* | | | | Th | 71 | Female | 25.3 | No | No | No | No | No | No | No | Right Hepatectomy | SCC | n/a | 152x114 | n/a | n/a | n/a | n/a | n/a |
| *Patient 22* | | | | Th | 58 | Female | 36.3 | No | No | No | No | No | No | No | Left Hepatectomy and portal lymphadenectomy | iCCA | n/a | 70x60 | Moderate | 1b | 1 | No | Yes |
| *Patient 23* | | | | Th | 71 | Male | 24.3 | No | Yes | No | No | No | No | No | Central Hepatectomy | HCC | n/a | 31x27 | Moderate | 1b | x | No | No |
| *Patient 24* | Ti | | P (ti) | Th | 80 | Male | 24.1 | No | No | No | No | No | No | No | Segment VI Resection | iCCA | n/a | 39x38 | Moderate | 1a | x | No | No |
| *Patient 25* | | | | Th | 62 | Female | 27.2 | No | No | No | No | No | No | No | Left Hepatectomy | iCCA | n/a | 72x32 | Poor | 1b | x | No | No |

**Supplementary Table 3. Patient demographics, clinical characteristics, operative details and histological features of tissue samples included across experimental cohorts.** Patient demographics, clinical characteristics, operative details, histological features and experimental use are shown for tissue donors included across the study. Experimental use is indicated as Ti, timepoint analysis; P (lib), spectral library generation; P (ti), temporal proteomics library generation; and Th, therapeutic response experiments. BMI, body mass index; B–C, Bismuth–Corlette classification; ETOH xs, excess alcohol intake; HCC, hepatocellular carcinoma; iCCA, intrahepatic cholangiocarcinoma; LVI, lymphovascular invasion; pCCA, perihilar cholangiocarcinoma; PNI, perineural invasion; PSC, primary sclerosing cholangitis; PVE, portal vein embolisation; SCC, squamous cell carcinoma. n/a, not applicable or not available.

| *Subtype* | *Day 0* | *Day 3* | *Day 7* | *Day 11* | *Day 15* |
|---|---|---|---|---|---|
| *pCCA* | 1.142 ± 0.812 (n = 9) | 1.328 ± 0.909 (n = 9) | 1.260 ± 1.073 (n = 9) | 0.731 ± 0.827 (n = 7) | 1.155 ± 0.765 (n = 6) |
| *iCCA* | 2.018 ± 0.855 (n = 8) | 1.915 ± 0.833 (n = 8) | 2.368 ± 1.116 (n = 8) | 1.627 ± 0.952 (n = 6) | 1.019 ± 0.946 (n = 7) |

**Supplementary Table 4. Viability differs between anatomical cholangiocarcinoma subtypes at baseline but remains measurable throughout culture.** Raw MTS absorbance values are presented as mean ± SD, with the number of patients contributing at each timepoint shown in parentheses. Day 0 MTS absorbance was higher in iCCA than pCCA tissue slices (Welch's two-sided t-test, $p = 0.048$); later timepoints are shown descriptively.

iCCA, intrahepatic cholangiocarcinoma; pCCA, perihilar cholangiocarcinoma.

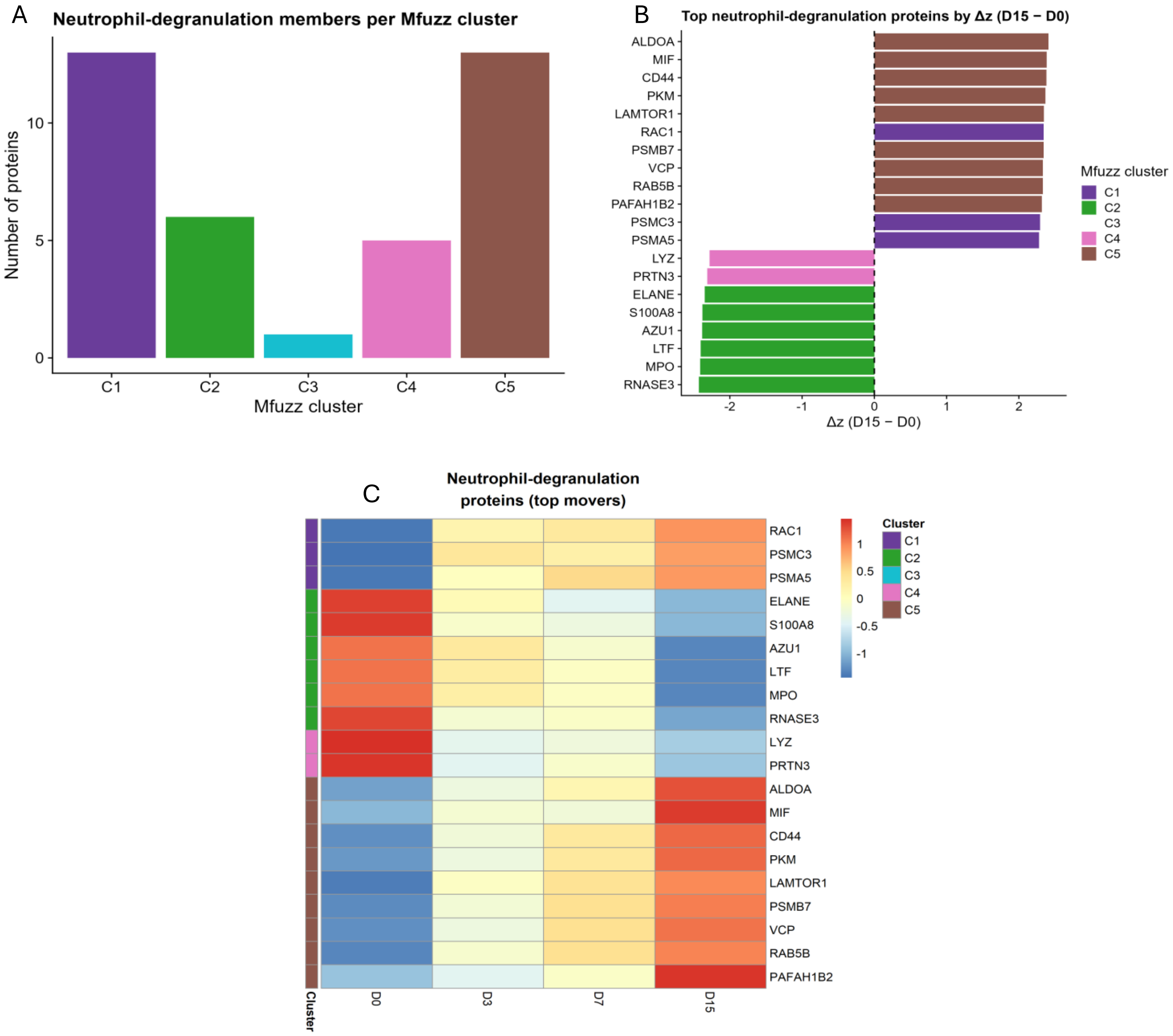


**Supplementary Figure 1. Neutrophil degranulation-associated proteins occupy distinct temporal programmes during tissue slice culture.** (A) Number of Reactome neutrophil degranulation proteins assigned to each Mfuzz cluster (C1–C5). (B) Top 20 neutrophil degranulation proteins ranked by absolute change in row z-score between Day 0 and Day 15 [Δz(D15–D0)]; positive values indicate increased and negative values decreased abundance over culture. (C) Row z-score heatmap showing temporal profiles of the same proteins across Days 0, 3, 7 and 15, grouped by Mfuzz cluster.

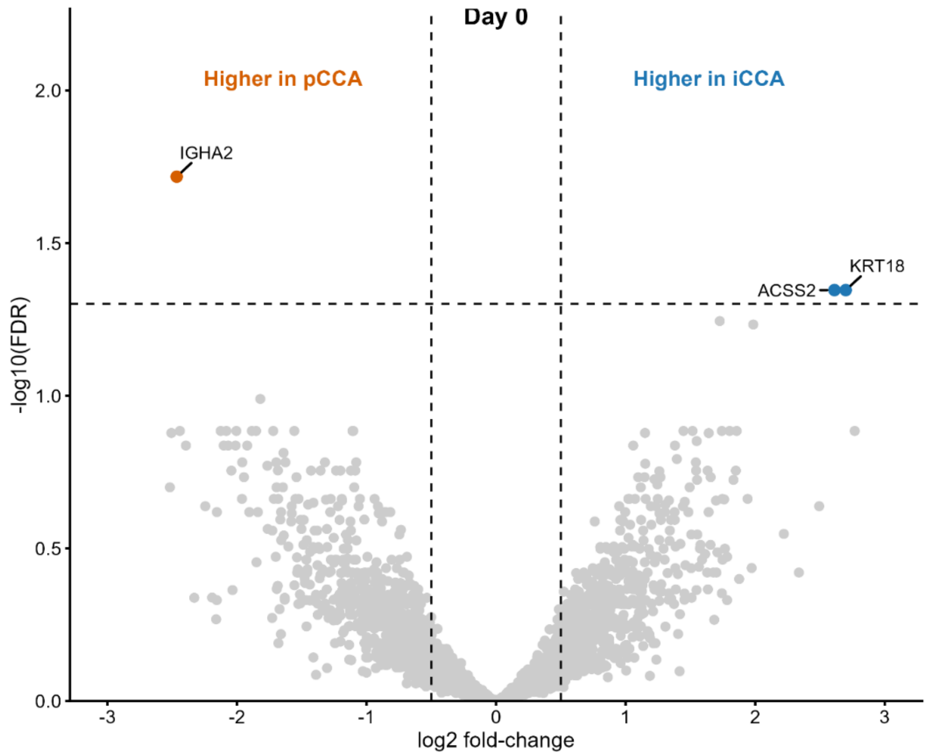


**Supplementary Figure 2. Baseline proteomic differences between cholangiocarcinoma subtypes are limited to a small number of proteins.** Volcano plot comparing protein abundance between iCCA and pCCA at Day 0 following detection filtering (3,750 of 4,578 protein groups). Positive log2 fold-change indicates higher abundance in iCCA and negative values higher abundance in pCCA. Blue and orange indicate proteins meeting FDR ≤ 0.05 and |log2 fold-change| ≥ 0.5, while grey indicates non-significant proteins. Dashed vertical lines indicate the ±0.5 log2 fold-change threshold and the horizontal dashed line indicates FDR = 0.05.

FDR, false discovery rate; iCCA, intrahepatic cholangiocarcinoma; pCCA, perihilar cholangiocarcinoma.
